# Manually weighted taxonomy classifiers improve species-specific rumen microbiome analysis compared to unweighted or average weighted taxonomy classifiers

**DOI:** 10.1101/2025.03.12.642789

**Authors:** Ryukseok Kang, Zhongtang Yu, Tansol Park

## Abstract

Previous research has demonstrated that applying taxonomic weights to shotgun metagenomic data can improve species identification in 16S rRNA gene-based microbiome analysis. However, such an approach does not allow for accurate analysis of samples collected from less studied habitats, such as rumen. In the present study, we developed a method to incorporate taxonomic weights based on relative abundance of species identified from shotgun sequencing and amplicon sequencing data derived from rumen. Using this weighting method, we evaluated latest versions of five prominent databases—SILVA, Greengenes2, RDP, NCBI RefSeq, and GTDB—against the BLAST 16S rRNA database, assessing classification counts, fully classified ratios (proportion of ASVs classified to a known genus and species), and error rates. Our results revealed that the use of the weighting method significantly improved both classification counts and fully classified ratios, along with a substantial (*P* < 0.05) reduction in error rates compared to unweighted taxonomy classifier. While GG2 and SILVA struggled with accurate classification at the species level owing to their inherent database characteristics, GTDB consistently improved all metrics using the manually weighted taxonomy classifier, achieving up to an 8% error rate reduction at the species level. NCBI RefSeq and RDP also exhibited remarkable improvement in the classification counts and fully classified ratios, along with substantial error rate reductions by up to 47% at the species level. These findings demonstrate that amplicon sequencing datasets can enhance rumen microbiome analyses through effective weighting methods. While SILVA is commonly used in metataxonomic analyses of the rumen microbiome, we recommend NCBI RefSeq for species-level classification due to its superior accuracy and minimal ambiguous classification (e.g., “uncultured” or “sp.") in future metataxonomic studies.

## Background

Ruminants host a diverse and highly specialized microbiome in their rumen, which enables them to digest a wide range of feedstuffs, particularly fibrous materials that would otherwise be indigestible [1, 2]. Accurately identifying the key microbes and understanding their contributions to various rumen fermentation processes are crucial for improving animal nutrition and productivity and mitigating the environmental impact associated with this unique animal group. The rumen microbiome has been intensively studied using culture-based and culture-independent nucleic acid-based techniques, with recent efforts predominantly leveraging the latter that utilize sequencing approaches like metataxonomics, metagenomics, and metatranscriptomics. Advances in sequencing technologies and the expansion of reference databases have significantly improved the accuracy of microbiome characterization, including taxonomic classification. However, most gut microbiome databases are disproportionally dominated by human microbiome data [3]. Such database bias may reduce the accuracy of analyses for non-human microbiomes, including the rumen microbiome.

To address this challenge, several initiatives have been proposed, including the Global Rumen Census project, the “Hungate1000,” “Holoruminant,” and meta-omic approaches aimed at enhancing comprehensive characterization of the rumen microbiome [4–7]. Despite these efforts, the accuracy of 16S rRNA gene-based metataxonomic analysis of the rumen microbiome remains uncertain. Recent advances, such as weighted taxonomy classifiers, have shown promise in improving taxonomic resolution. For example, Kaehler et al. (2019) utilized q2-clawback and the Earth Microbiome Project Ontology (EMPO) 3 datasets to apply weighting to taxonomy classifiers. Specifically, this approach assigns varying weights to individual taxa: higher weights to taxa prevalent in specific environments and lower weights to rare or absent taxa [8]. This approach improved species-level identification and abundance estimation accuracy. It was also suggested that weighted taxonomy classifiers could further improve taxonomic resolution when applied to metagenomic analyses. However, since the ruminal microbiome is not included in the EMPO habitat types, the proposed classifiers probably cannot be applied to analyses of rumen samples.

The distinct rumen microbiome profiles observed across ruminant species [9, 10] suggest that generalized taxonomic weights may compromise classification accuracy when applied to a specific ruminant species. Therefore, creating and implementing ruminant species-specific taxonomy classifiers would be essential for achieving greater classification accuracy. Additionally, while shotgun sequencing-based metagenomics has been increasingly used for improved resolution, 16S rRNA gene amplicon-based metataxonomics remains widely used for rumen microbiome profiling due to its cost-effectiveness. To achieve greater accuracy in metataxonomic analyses of the rumen microbiome, we utilized multiple datasets of partial and full-length 16S rRNA gene sequences along with shotgun sequences obtained from native Korean Hanwoo cattle in developing weighted taxonomic classifiers. Furthermore, unlike the earlier studies by Kaehler et al. (2019) that used an outdated version of Greengenes for validating weighted methods, we used the most recent Greengenes database (GG2). We also included the latest versions of other commonly used databases—SILVA, NCBI RefSeq, GTDB, and RDP—to explore how the choice of databases affects classification accuracy. We developed an enhanced taxonomy classification method that integrates weight assignment, additional preprocessing steps, and q2-clawback by utilizing data from native Korean Hanwoo cattle. This study indicates that applying taxonomic weights can significantly improve rumen microbiome analyses.

## Methods

### Constructing an unweighted and weighted taxonomy classifier

All data were analyzed using the QIIME2-amplicon version 2024.10 [11]. To construct unweighted taxonomy classifier (UWTC), the NCBI reference sequence database (RefSeq), SILVA, and GTDB databases were collected through the q2-rescript [12]. The NCBI RefSeq was downloaded on October 14, 2024, and used for archaeal and bacterial 16S rRNA sequences. The SILVA versions 138.1 and 138.2 [13], GTDB version 220.0 [14], along with the GG2 versions 2022.10 and 2024.09 [15] and the RDP bacterial and archaeal hierarchy model (version 2.14, August 2023) [16], were used to build the taxonomy classifier. Archaeal sequences shorter than 900 bp and bacterial sequences shorter than 1,200 bp were filtered out from all the databases to enhance taxonomic reliability and resolution. Additionally, all the database sequences were dereplicated to eliminate redundancy.

To construct an average weighted taxonomy classifier (AWTC) as described in the QIIME2 resource (https://resources.qiime2.org/), metadata were developed using the Earth Microbiome Project Ontology (EMPO) [17] for assigning average taxonomic weights. Publically available V4 region (150 bases) of the 16S rRNA gene sequences and associated EMPO3 metadata were obtained from Qiita using q2-clawback [8]. The following keywords were used to search and download the EMPO3 metadata: ‘animal-corpus,’ ‘animal-distal-gut,’ ‘animal-proximal-gut,’ ‘animal-secretion,’ ‘animal-surface,’ ‘plant-corpus,’ ‘plant-rhizosphere,’ ‘plant-surface,’ ‘sediment-non-saline,’ ‘sediment-saline,’ ‘soil-non-saline,’ ‘surface-saline,’ ‘water-non-saline,’ ‘water-saline,’ ‘animal-non-saline,’ and ‘animal-saline.’

Metagenomic shotgun and 16S rRNA gene amplicon datasets derived from rumen fluid samples collected directly from Hanwoo cattle were used to construct manually weighted taxonomy classifier (MWTC). The characteristics of each dataset are presented in Table 1. The metagenomic shotgun dataset from Hanwoo steers (36 samples) and cows (5 samples) was preprocessed using fastp [18] for quality control, followed by sequence filtering with Bowtie2 (version 2.5.4) [19] to remove host sequences by aligning them to the Hanwoo genome (GCA_028973685.2) and feed sequences by aligning them to the genomes of the feed ingredients consumed by each animal group. For Hanwoo steers, feed sequences were filtered out using the genomes of oat hay (GCA_916181665.1), corn (GCF_902167145.1), rice straw (GCF_001433935.1), wheat (GCF_018294505.1), and palm kernel meal (GCF_000442705.1). For cows, filtering was performed using oat hay, corn, and rice straw genomes. The filtering ensured the isolation of microbial DNA sequences by removing host and feed-related sequences. The filtered metagenomic datasets were processed using SortMeRNA (version 4.3.7) [20] to extract the rRNA gene sequences. The resultant rRNA sequences were directly imported into QIIME2, and a feature table was generated after sequence dereplication. For the 16S rRNA gene amplicon sequencing data, raw reads were imported directly into QIIME2, followed by merging the paired-end amplicon sequences using FLASH2 (version 2.2.00) [21] with a minimum overlap of 20 bp. The sequences were then denoised using Deblur [22] with default options.

**Table 1.** Summary of the datasets used to generate the manual weighted taxonomy classifier.

| NCBI<br>BioProject | Platform | Number<br>of<br>samples | Number<br>of<br>sequences<br>(mean ± SD) | References |
| --- | --- | --- | --- | --- |
| - | Illumina HiSeq | 36 | 363,025 ± 89,416 | Unpublished |
| - | Illumina HiSeq | 5 | 388,852 ± 79,987 | Unpublished |
| - | PacBio Sequel | 15 | 81,209 ± 8,773 | [47] |
| PRJEB25166 | Illumina MiSeq | 9 | 30,085 ± 2,475 | [48] |
| PRJNA523867 | Illumina MiSeq | 25 | 54,052 ± 4,803 | [49] |
| PRJNA725944 | Illumina MiSeq | 8 | 47,213 ± 13,503 | [50] |
| PRJNA797687 | Illumina MiSeq | 184 | 33,261 ± 16,056 | Unpublished |

A custom Python script was used to combine the feature tables and the corresponding representative sequences to generate the input file for taxonomy weight generation. Sequences associated with chloroplasts and mitochondria were removed before generating the classifier to ensure accuracy. Weighted datasets were then generated using q2-clawback, utilizing 16S rRNA gene sequences from the EMPO dataset. These datasets incorporated all databases used to create the classifier, including UWTC generated with the naïve Bayes algorithm using the ‘feature-classifier’ plugin [23]. The process for constructing the weighted taxonomy classifier is detailed in Fig. 1.

**Fig. 1.**
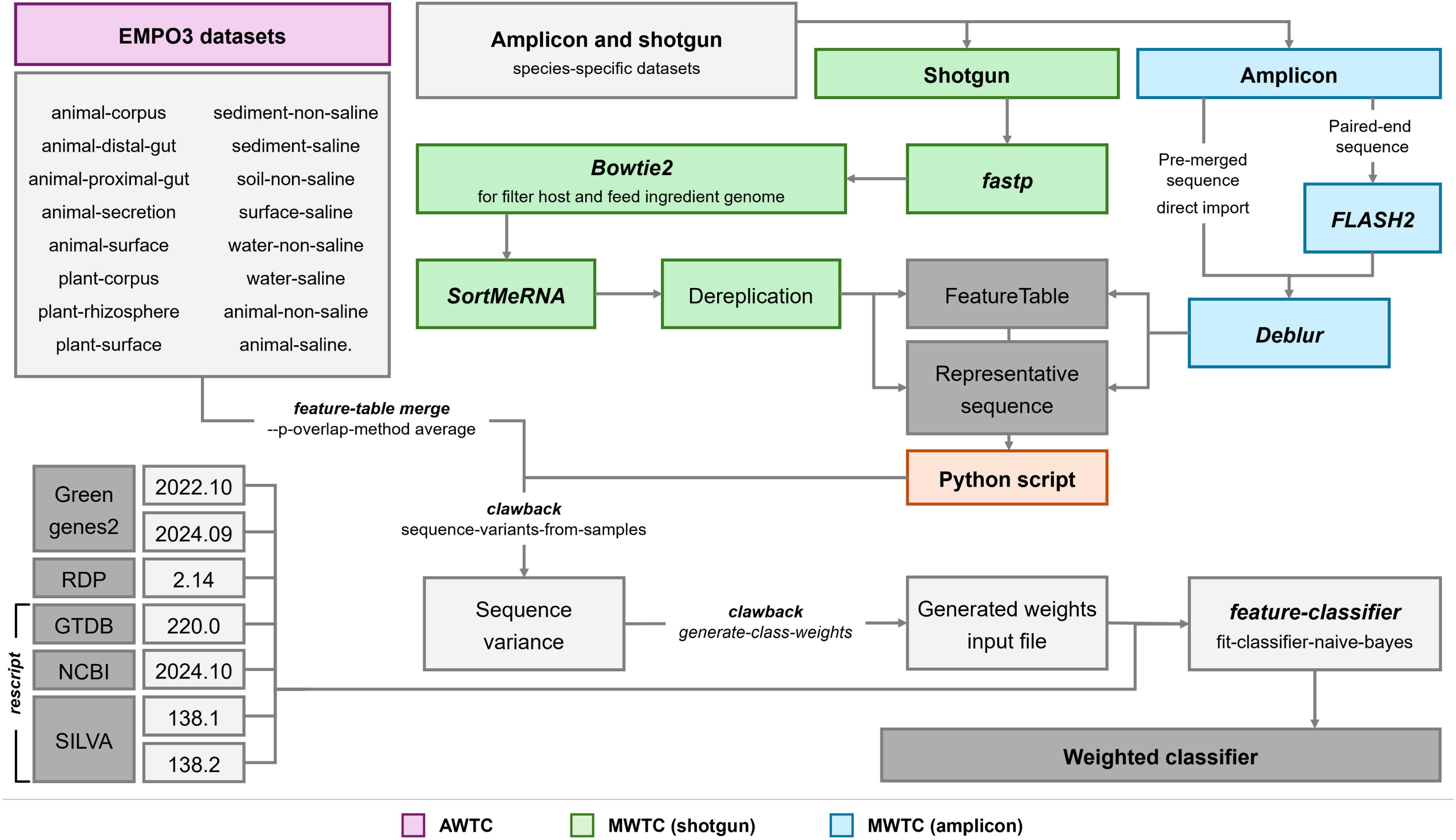
The workflow used to construct each weighted taxonomy classifier. The bioinformatics tools used were italicized.

### Amplicon datasets for validation of taxonomy classifier

16S rRNA gene amplicon sequencing data were generated from rumen samples of Hanwoo cattle to validate the taxonomy classifiers. Briefly, rumen fluid samples were directly collected via stomach tubing from the rumen following an *in vivo* experiment and *in vitro* fermentation for 24 h. The nearly full-length and the V3-V4 hypervariable region of the 16S rRNA gene were amplified using the primers 27F (5′-AGRGTTYGATYMTGGCTCAG-3′) and 1492R (5′-GYTACCTTGTTACGACTT-3′) and 341F (5′-CCTACGGGNGGCWGCAG-3′) and 805R (5′-GACTACHVGGGTATCTAATCC-3′), producing amplicons of full-length and V3-V4 regions, respectively. A total of 47 *in vivo* samples and five *in vitro* samples were used for validating the classifiers with the full-length amplicons, while 30 *in vivo* samples and four *in vitro* samples were analyzed for the V3-V4 amplicons.

### Data processing

The amplicon sequences were classified into 21 classifiers using scikit-learn [24], employing three classification methods—UWTC, AWTC, and MWTC—applied across the five databases. For comparisons of the overall microbiota composition between the databases and among UWTC, AWTC, and MWTC, principal component analysis (PCA) was conducted using Bray-Curtis dissimilarity matrices at both the genus and species levels for both the full-length and the V3-V4 amplicon sequences. Alpha diversity metrics, including observed features [25], evenness [26], Shannon [27], Simpson, and inversed Simpson diversity indices [28] were also computed at these levels. Prior to these analyses, the data from all samples were normalized using Total Sum Scaling (TSS). Classification counts were calculated for all amplicon sequence variants (ASVs) only when there was no blank classification at each taxonomic level, from the phylum to the species. To address cases where certain ASVs were not fully classified at a specific taxonomic level, the ratio of fully classified taxa was calculated as the number of fully classified taxa at each level relative to the total taxa count. This metric, referred to as the fully classified ratio, was evaluated for each database at both the genus and species levels to assess classification efficiency.

### Error rate estimation

Error rates for taxonomic classification were calculated following the approach outlined by Kaehler et al. [8] to determine the proportion of ASVs misclassified by classifiers. The BLAST 16S ribosomal RNA database [29] was used as a reference, and all ASVs were classified to the species level without considering confidence scores. This ensured that every ASV was classified to a species regardless of classification confidence. However, significant updates in taxonomy occurred after the release of GG2 2022.10 and SILVA 138.1, and in these updates some taxa were reclassified at both the phylum (e.g., *Firmicutes* to *Bacillota*, *Bacteroidetes* to *Bacteroidota*, *Proteobacteria* to *Pseudomonadota*, and *Actinobacteria* to *Actinomycetota*) and the genus levels (e.g., certain *Prevotella* species reclassified to *Xylanibacter* and *Segatella*, and *Propionibacterium acnes* renamed as *Cutibacterium acnes*) [30, 31]. Thus, error rates were not calculated for these databases.

### Statistical analysis

The effects of classifier types on overall microbiota composition analysis were statistically evaluated using permutational multivariate analysis of variance (PERMANOVA). This analysis used the vegan and pairwiseAdonis packages in R (version 4.3.3) with 9,999 random permutations [32, 33]. Multiple testing adjustments were applied using the Benjamini-Hochberg correction method [34]. Normality of alpha diversity metrics, classification counts, fully classified ratios, and error rates were assessed using the Shapiro-Wilk test, while homogeneity of variances was evaluated using Levene’s test. All data satisfied the assumption of homogeneity of variances. Statistical analysis was conducted using PROC MIXED in SAS 9.4 (SAS Institute Inc., Cary, NC, USA) for normally distributed data, whereas PROC GLIMMIX was used for the data that did not follow a normal distribution. Classifier type was treated as a fixed effect. For the alpha diversity metrics, classification counts and fully classified ratios, each amplicon dataset was included as a random effect, while for the error rates, both amplicon dataset and database types were treated as random effects. Statistical significance was declared at s threshold of *P* ≤ 0.05.

## Results

The taxonomy classifier evaluation method was validated using microbiota datasets derived from both *in vitro* experiments and *in vivo* samples collected from the rumen of Hanwoo cows. The *in vitro* fermentation experiment involved the collection of ruminal fluid and incubation with a mixed microbial substrate under controlled conditions, enabling the study of fermentation dynamics and microbial composition. This approach is widely used in ruminant research [35]. A recent study validated this approach by demonstrating that *in vitro* batch cultures of rumen fluid effectively maintained the ruminal microbiome for up to 48 h [36]. Based on this evidence, both *in vivo* and *in vitro* datasets were selected to validate the taxonomy classifiers.

### Classifier evaluation using in vivo datasets

The classifiers were comparatively evaluated for diversity analyses and taxonomic identification of rumen microbes. To compare within-sample diversity and evaluate different classifiers, alpha diversity analysis was performed at the genus and species level (Tables S1 and S2). Observed features were statistically different (*P* < 0.05) across all classifiers and taxonomy levels, except for GTDB with full-length amplicon sequences at the species level (*P* = 0.0526). The MWTC exhibited the highest values of observed features compared to the UWTC and AWTC with NCBI RefSeq and RDP (*P* < 0.01) but either lower or comparable diversity metric values with GG2, SILVA, and GTDB (*P* < 0.01). With the full-length amplicon sequences, MWTC showed the lowest evenness, Shannon index, and Simpson index compared to UWTC with NCBI RefSeq and RDP (*P* < 0.01), but with the V3-V4 amplicon sequences, these alpha diversity metrics were higher or comparable. When GTDB was used, AWTC and MWTC yielded higher Shannon, Simpson, and Inverse Simpson indices at the genus level compared to UWTC (*P* < 0.01) but lower indices at the species level (*P* < 0.01).

To assess differences in microbiota identified by the three taxonomy classifiers (i.e., UWTC, AWTC, and MWTC), we performed beta diversity analysis using PERMANOVA. Significant differences (*P* = 0.001) were noted in microbiota composition depicted by the full-length amplicon sequences at the genus level with NCBI RefSeq and RDP, while no significant differences were noted with the other databases (Fig. 2A). At the species level, however, the revealed microbiota compositions were significantly affected (*P* = 0.001) by the taxonomy classifier used irrespective of databases (Fig. 2B). With respect to the microbiota composition revealed by the V3-V4 amplicon sequences, significant differences were identified at both the genus and the species levels across all databases (*P* = 0.001) (Fig. 2C, D). Pairwise comparisons revealed that with the GG2, microbiota profiles classified using UWTC and MWTC were similar, whereas AWTC produced distinct microbiota profiles. Significant differences in microbiota profiles were observed among UWTC, AWTC, and MWTC with all other databases (Table S3).

**Fig. 2.**
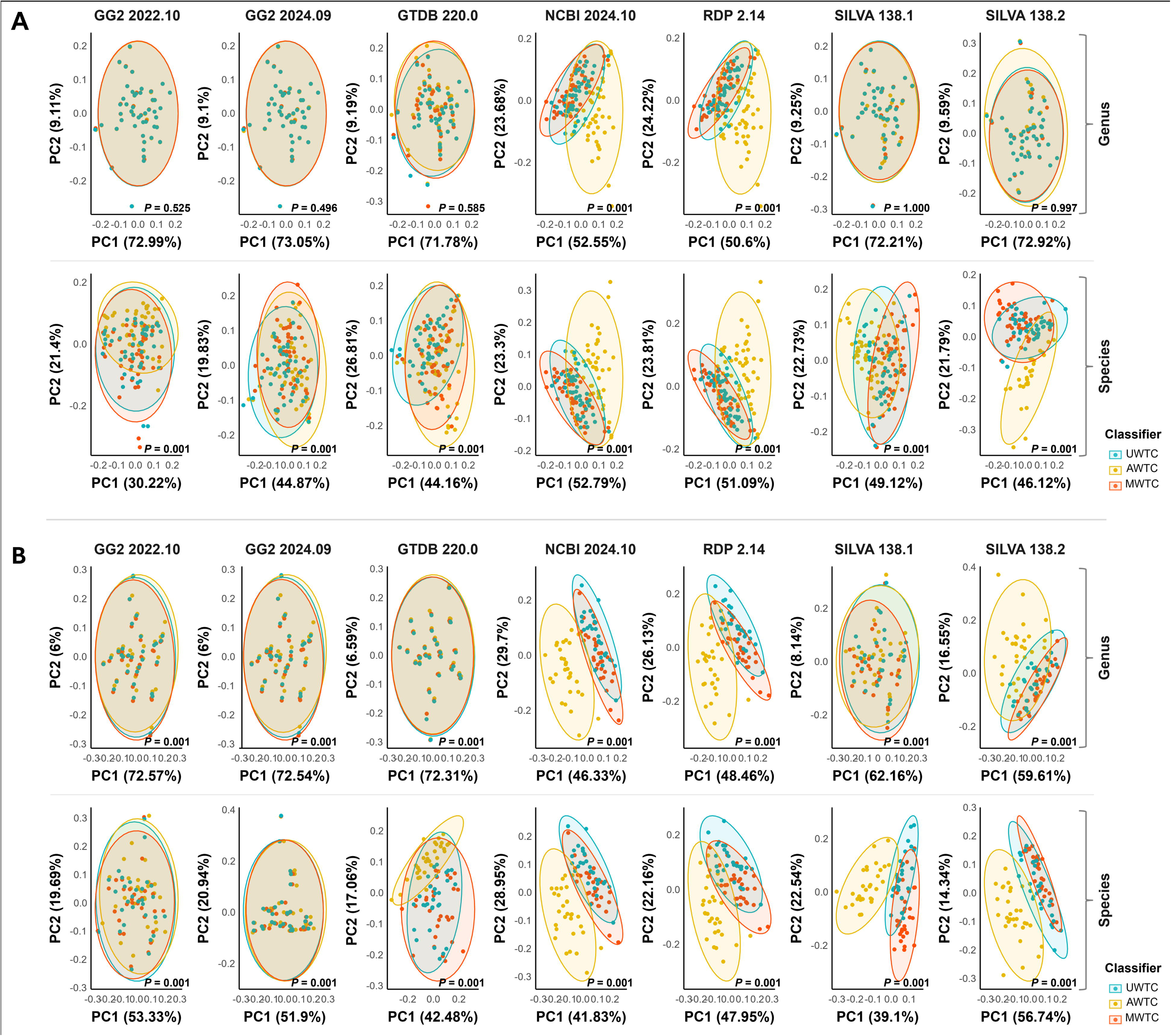
PCA plots showing differences in rumen microbiomes among taxonomy classifier types within each database at genus and species levels for full-length amplicon sequences (A) and for V3-V4 amplicon sequences (B) in an *in vivo* study. Significant differences were observed for all classifiers except for NCBI RefSeq and RDP at the genus level.

The classifiers were evaluated for taxonomic classification of key rumen microbial taxa and the proportion of successfully annotated taxa, focusing on the top 20 most dominant genera and species. Across all databases, MWTC yielded higher proportion of the top 20 dominant taxa compared to UWTC and AWTC, while maintaining the same taxonomic composition (Fig. 3, S2). With NCBI RefSeq and RDP, AWTC assigned some of the full-length and V3-V4 amplicon sequences to *Segatella copri*, whereas MWTC and UWTC assigned these sequences to *Xylanibacter ruminicola* (Fig. 3A, S2B). With other databases, UWTC and MWTC identified similar dominant taxa. With GG2 and GTDB, over 70% of the top 20 taxa were classified as "sp.," whereas with SILVA, more than 80% of the top 20 taxa were annotated as "uncultured." Notably, with SILVA at the species level, AWTC predominantly assigned uncultured microorganisms as "bacterium," whereas MWTC annotated them as "rumen."

**Fig. 3.**
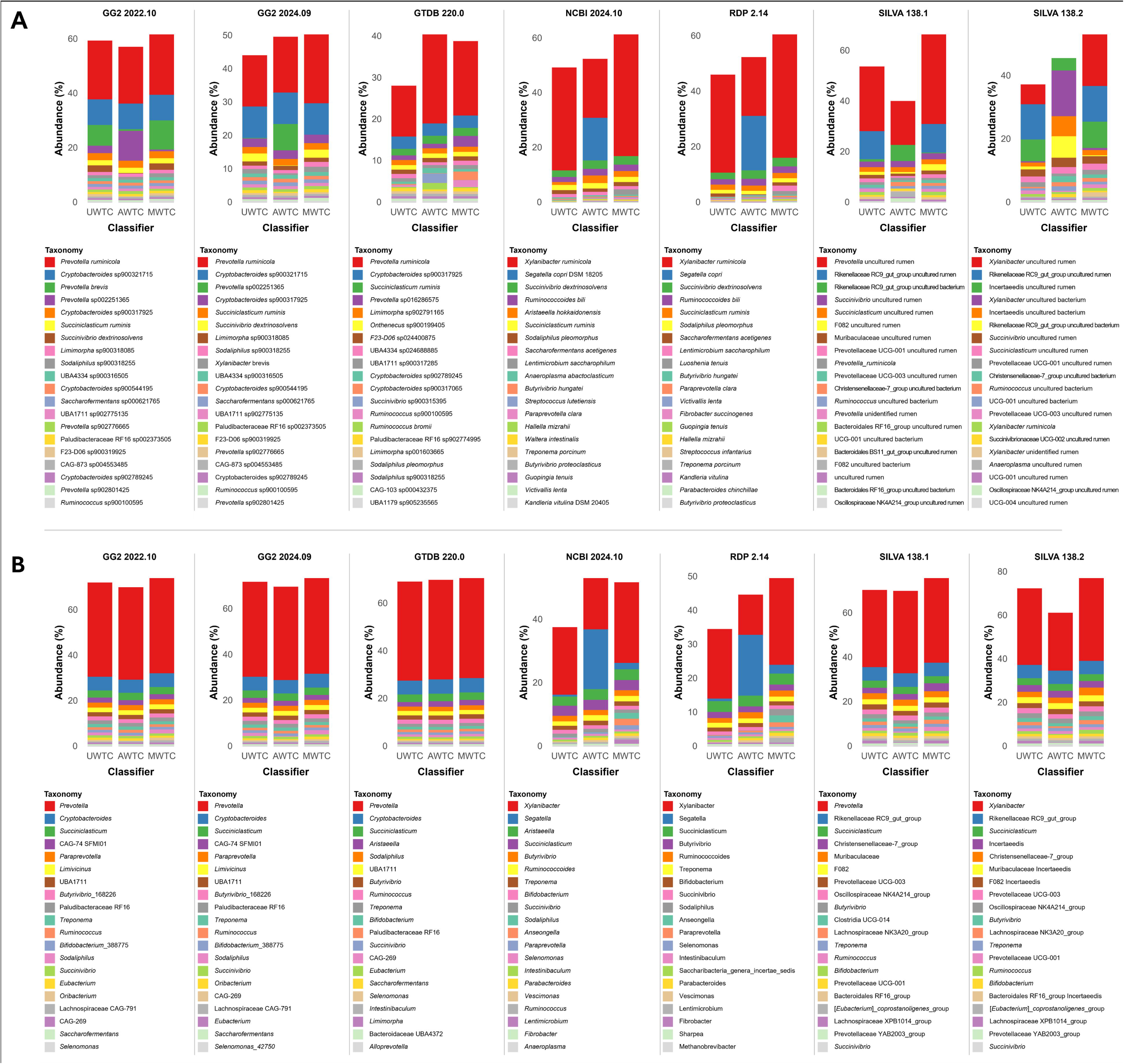
Taxonomic barplot showing the top 20 taxa for each taxonomy classifier in an *in vivo* study. Taxonomy classification was performed at the species level using full-length amplicon sequences (A) and at the genus level using V3-V4 amplicon sequences (B).

In terms of classification counts and fully classified ratios, MWTC outperformed UWTC and AWTC with both the full-length amplicon sequences and V3-V4 amplicon sequences across all databases (Fig. 4, 5). Specifically, MWTC exhibited similar or higher classification counts at the genus level although it showed lower or no difference in classification counts at the phylum and family levels compared to UWTC and AWTC with the GG2 and SILVA databases. At the species level, MWTC increased classification counts by over 5% and 40% with the GG2 and the SILVA databases, respectively, compared with UWTC (*P* < 0.001). Similarly, with NCBI RefSeq and RDP, MWTC demonstrated a >20% increase in higher classification counts and fully classified ratios over UWTC and even greater increases over AWTC at both the genus and species levels (*P* < 0.001) (Tables S2 and S3). Notably, with SILVA 138.1 version, AWTC performed worse than UWTC with the full-length amplicon sequences. With the V3-V4 amplicon sequences, AWTC also performed worse than UWTC at the genus level but slightly better than MWTC at the species level (Tables S4 and S5).

**Fig. 4.**
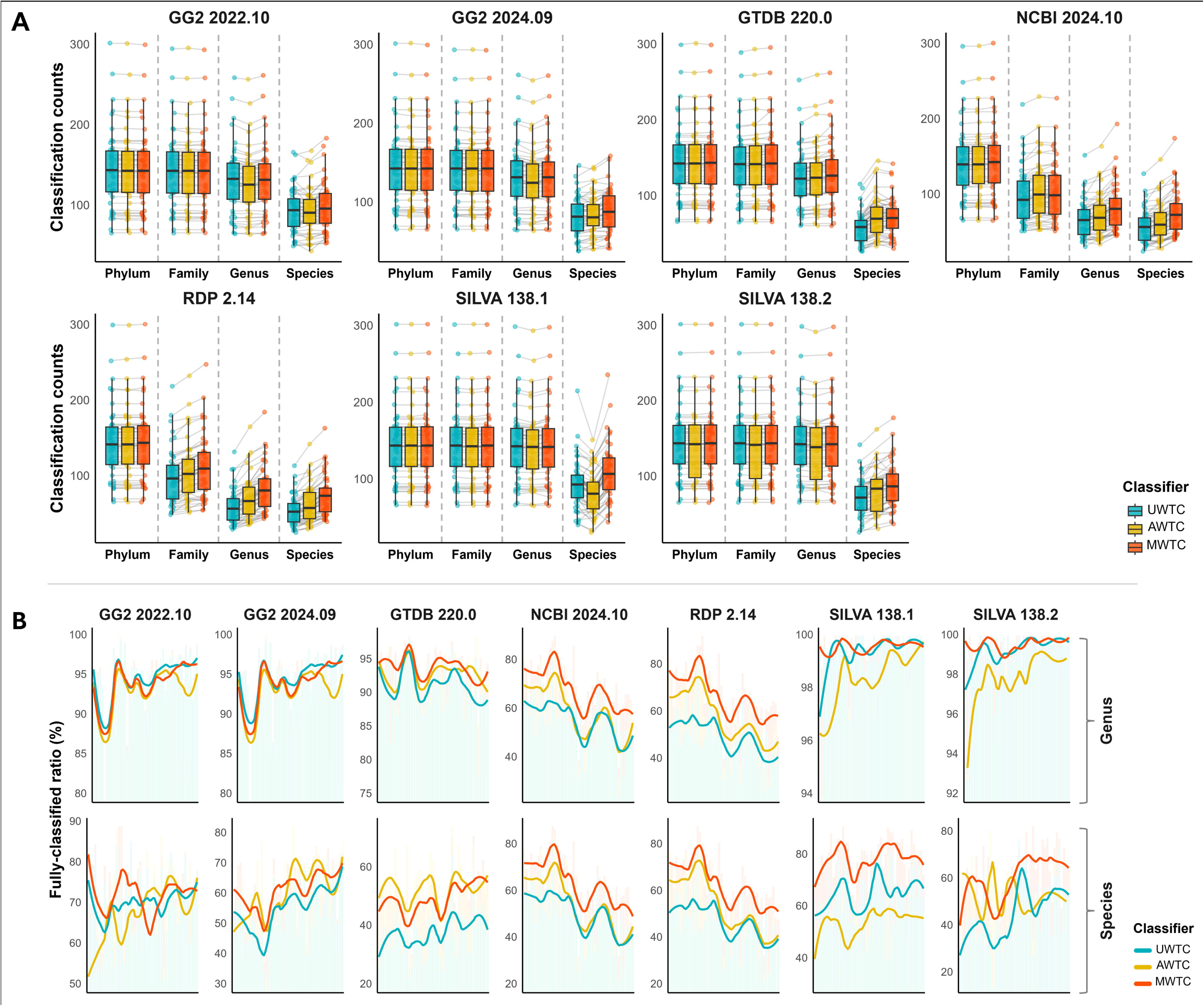
Counts of classified ASVs at the phylum, family, genus, and species levels for each database (A) and the fully classified ratios (the proportion of completely classified features relative to the total ASVs) (B) for each database identified using the full-length amplicon sequences in an *in vivo* study. Classifiers were categorized as follows: UWTC, a classifier without any adjustments; AWTC, a classifier based on average data from the EMPO3 dataset; and MWTC, a classifier manually curated using metagenomic and amplicon sequencing data.

**Fig. 5.**
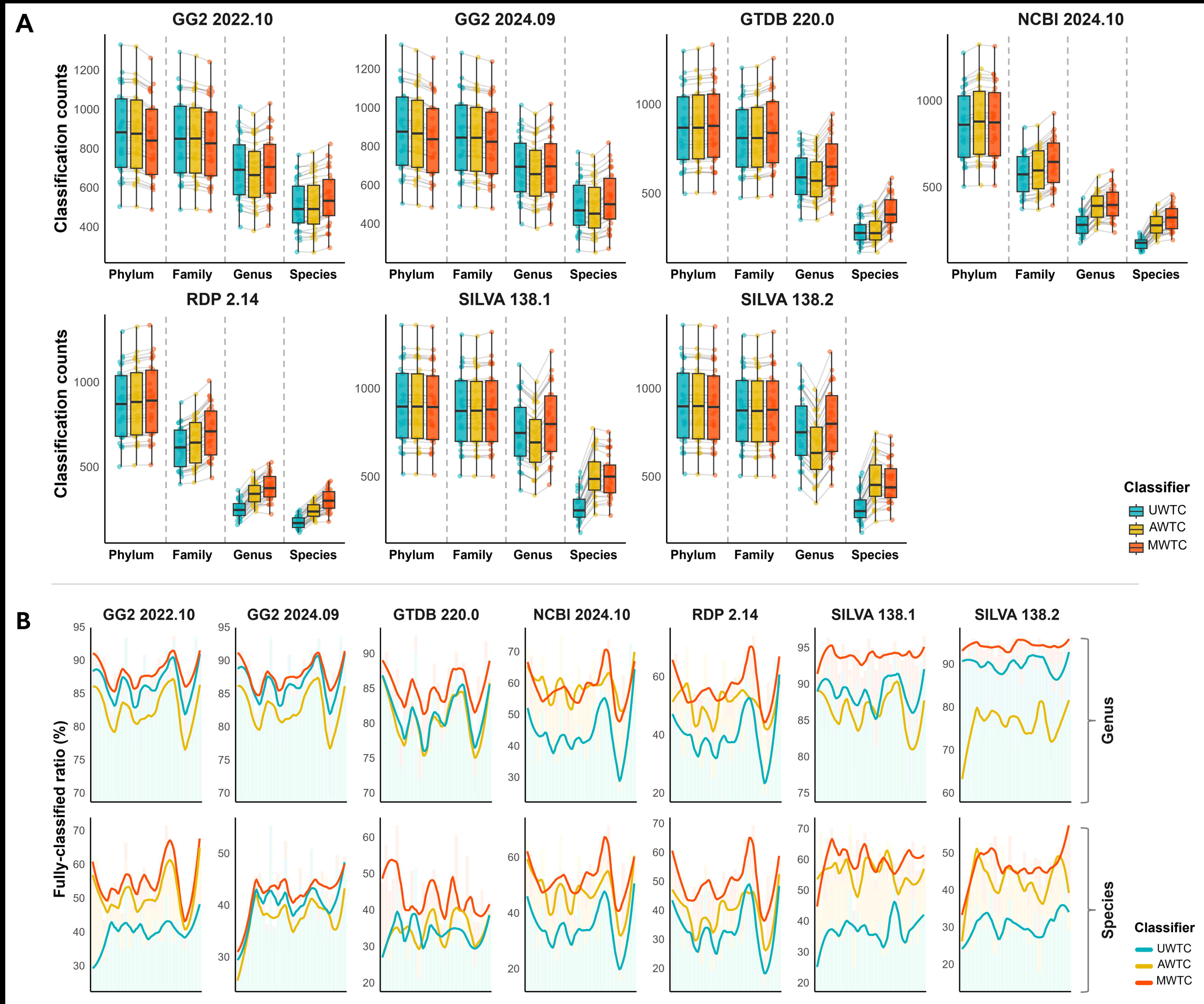
Counts of classified ASVs at the phylum, family, genus, and species levels for each database (A) and the fully classified ratios (the proportion of completely classified features relative to the total ASVs) (B) for each database identified using the V3-V4 amplicon sequences in an *in vivo* study. Classifiers were categorized as follows: UWTC, a classifier without any adjustments; AWTC, a classifier based on average data from the EMPO3 dataset; and MWTC, a classifier manually curated using metagenomic and amplicon sequencing data.

### Classifiers evaluation using in vitro experimental samples

*In vitro* experiments are often used to simulate the rumen environment when conducting preliminary comparisons or when large-scale animal trials are not feasible. Therefore, since 16S rRNA amplicon sequences derived from *in vitro* experiments also require improved resolution through MWTC, we validated MWTC using *in vitro* experimental samples.

The alpha diversity indices demonstrated notable differences across classifiers (Tables S6 and S7). Observed features were statistically significant across all classifiers (*P* < 0.05), except for GTDB with full-length amplicon sequences. Consistent with the *in vivo* datasets, NCBI RefSeq and RDP demonstrated higher observed features with MWTC compared to UWTC (*P* < 0.05), whereas GG2 and SILVA exhibited lower or similar values. Shannon, Simpson, and inverse Simpson indices were significant in both full-length and V3-V4 amplicon sequences, except for GG2 and SILVA at the genus level in full-length sequences (*P* < 0.05). When used with NCBI RefSeq and RDP, MWTC yielded higher values in these indices compared to UWTC and AWTC, irrespective of sequencing types or taxonomic levels. In contrast, coupled with GG2 and SILVA, MWTC exhibited lower or similar values than UWTC. AWTC displayed inconsistent trends, with the values fluctuating, relative to UWTC.

The beta diversity analysis revealed no significant differences as determined with the full-length amplicon sequences at the genus level with the two SILVA database releases (Fig. S1A). In contrast, other genus-level data, except for the SILVA databases, and all species-level data were statistically different in beta diversity (*P* < 0.05) (Fig. S1A, B). Both genus- and species-level data determined by the V3-V4 amplicon sequences differed statistically significantly across all databases (Fig. S1C, D), consistent with the *in vivo* results (Table S8).

Similar to the *in vivo* dataset, MWTC achieved the highest proportion accounted for by the top 20 dominant taxa compared to UWTC (Fig. S3, S4). MWTC showed an overall higher microbial abundance than UWTC. Across all the classifiers, at the species level, with GG2 and GTDB, 75% of the top 20 taxa were classified as "sp.," whereas with SILVA, 90% of the top 20 taxa were annotated as "uncultured" or "unidentified."

Regarding classification counts and fully classified ratios, MWTC consistently outperformed UWTC at both the genus and species levels (Fig. S5, S6). With the V3-V4 amplicon sequences, AWTC and MWTC yielded lower classification counts than UWTC at the phylum and family levels when the GG2 and SILVA databases were used. Similarly, with the full-length sequences and the V3-V4 amplicon sequences, AWTC showed lower values in classification counts than UWTC at the genus and species levels when GG2 was used, with a 3.4–5.5% reduction at the genus level and a 5.6–9.5% reduction at the species level. Similarly, when used together with SILVA, AWTC showed a 0.9–9.8% decrease in classification counts at the genus level. In contrast, while MWTC demonstrated lower values in classification counts than UWTC, these were statistically comparable to UWTC (Tables S9 and S10). Interestingly, with the NCBI RefSeq and RDP, MWTC exhibited fully classified ratio at the species level, with improvements ranging from 19.2–64.6% in NCBI RefSeq and 22.8–58.3% in RDP, which exceeded that achieved by UWTC.

### Error rate estimate

The error rates were thoroughly analyzed, taking into account variations across all databases for each amplicon dataset resulting from both *in vivo* and *in vitro* environments (Fig. 6). With the NCBI RefSeq, MWTC showed a relatively decreased in error rate compared to UWTC, averaging by 33.9–62.6% at the genus level and 22.6–47.5% at the species level. Similarly, with RDP, reductions of 25.2–53.5% and 15.7–24.9% were observed at the genus and species levels, respectively. The GTDB database yielded a decrease of 1.7– 4.4% at the genus level and 2.9–8.0% at the species level. With GG2 and SILVA, reductions were minimal, with a maximum decrease of only 1.2% at the species level. Overall, MWTC exhibited lower error rates than UWTC (*P* < 0.05), except at the phylum level with the V3-V4 amplicon sequences. Notably, the average error rate at the genus and species levels was higher with AWTC than with UWTC but lower with MWTC (Table S11).

**Fig. 6.**
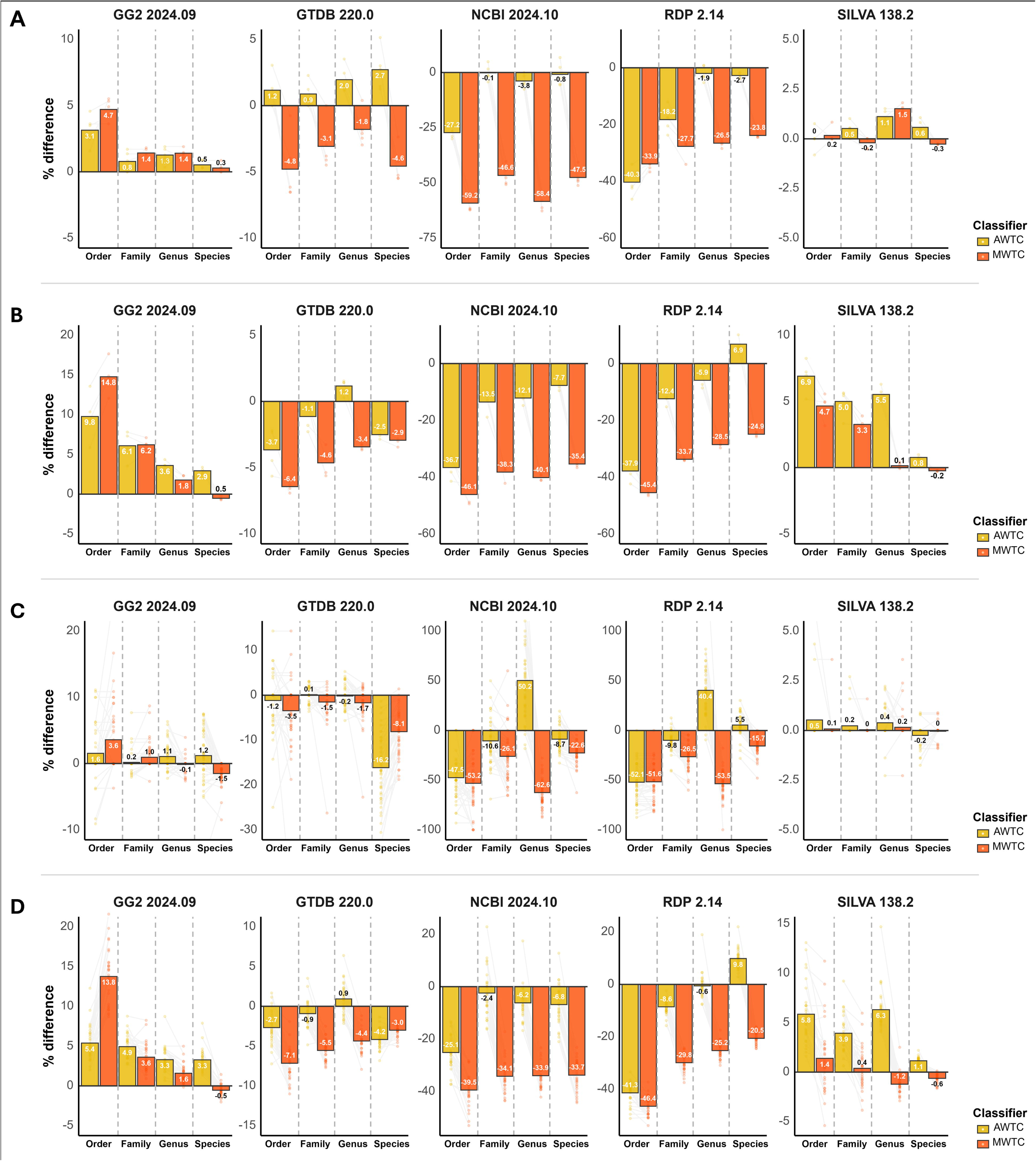
Error rate plots for each classification count, including *in vivo* experimental datasets from full-length amplicon sequences (A) and the V3-V4 amplicon sequences (B) and *in vitro* datasets from the full-length amplicon sequences (C) and the V3-V4 amplicon sequences (D). Error rates were calculated by comparing the results with those of the BLAST 16S ribosomal RNA database, considering both unnamed and misnamed classifications as errors.

## Discussion

Advancements in sequencing technologies and taxonomy classifiers have improved microbiome characterization, yet existing databases remain heavily human-centric, limiting analysis accuracies of non-human microbiomes like the rumen. While weighted taxonomy classifiers enhance microbial identification, their application to rumen samples may be less precise due to the absence of ruminal microbiota in EMPO habitat types and the unique microbial profiles influenced by ruminant species and diet. This underscores the need for ruminant-specific taxonomy classifiers to improve classification accuracy. To enhance the accuracy of taxonomic classification in rumen microbiome analyses, we suggested assigning weights during the classification process. However, since many microbiome databases do not contain sufficient rumen-specific data, we proposed the development of manually weighted datasets using both shotgun metagenomics and amplicon sequencing data from specific ruminant breeds.

### The MWTC constructed by integrating both shotgun and amplicon datasets could enhance taxonomic resolution

MWTC achieved higher classification counts, fully classified ratios, and a lower error rate than UWTC, demonstrating its capability to provide more accurate annotations, particularly at lower taxonomic levels. This suggests that even with the same ASV dataset, beta diversity analysis results could vary depending on the classifier used, which assigns different taxonomy. This further highlighted that classifiers with different taxonomy weighting methods influence the microbial community profiles differently.

MWTC preserves or increases the relative abundance of key microbial taxa present in the rumen samples. With NCBI RefSeq and RDP, AWTC reclassified *Xylanibacter ruminicola*, classified by UWTC, as *Segatella copri*. *Segatella copri* accounts for approximately 30% of the human gut microbiome [37, 38], but it is not dominant in the rumen. In contrast, MWTC retained the annotation of *Xylanibacter ruminicola* and further enhanced its relative abundance. Additionally, MWTC showed *Xylanibacter, Succiniclasticum, Succinivibrio, Sodaliphilus, Ruminococcoides, Butyrivibrio, Treponema,* and *Fibrobacter*, all of which are common genera found in the rumen [9, 39, 40], without a decline in abundance compared to UWTC. These suggest that MWTC provides a more accurate classification for rumen microbiota compared to the weighting approach used in AWTC.

A previous study had shown that using shotgun metagenome data as a weighting factor increased the taxonomy detection rate [8]. In this study, by incorporating not only shotgun sequencing datasets but also amplicon sequencing datasets as taxonomic weights, MWTC demonstrated higher classification counts and fully classified ratios, along with reduced error rates, than UWTC and AWTC. This indicated that amplicon sequencing datasets can be utilized to provide taxonomic weights while maintaining high resolution. However, not all the amplicon datasets are suitable for this purpose. To construct the MWTC, amplicon datasets derived from DNA collected directly from the ruminal fluid of individual animals were utilized. In contrast, amplicon datasets obtained from the *in vitro* fermentation process may not be ideal for use as weighting datasets, since they originate from mixed cultures gathered from multiple individuals [41, 42] and may include microorganisms introduced during the process of preparing buffer solutions [43, 44].

### The selection of taxonomy classifier databases is crucial at lower taxonomic levels

When GG2 and SILVA were used, no differences in beta diversity or error rate were noted. This finding could be attributed to the similar taxonomies implemented in GG2 and SILVA. In GG2, ambiguous annotations, such as ‘CAG,’ ‘RUG (Rumen uncultured genus)’ and ‘sp.’ are frequently observed, while in SILVA, annotations like ‘uncultured,’ ‘unidentified,’ or ‘unknown organism’ are common, both presenting challenges in identifying microbes at lower taxonomic levels and compromising taxonomic and phylogenetic rigor (Supplementary data) [12, 45]. Despite AWTC and MWTC showed higher classification count and fully classified ratio compared to UWTC, the error rate did not decrease in the GG2 and SILVA databases. The AWTC showed increased error rates compared to the UWTC at both the genus and species levels. However, the MWTC demonstrated lower error rates than the AWTC and overall showed similar or lower error rates than UWTC. This suggests that the MWTC maintained a resolution comparable to that of the UWTC when GG2 and SILVA were used.

The GTDB, which serves as the underlying database for GG2, shares the same classification framework as GG2 [15]. Similar to GG2, GTDB reflects the characteristics of the database with annotations like ‘CAG,’ ‘RUG (Rumen uncultured genus)’ and ‘spp.’ However, unlike GG2 and SILVA, GTDB did not yield lower classification counts at the phylum or family levels and showed a slight improvement in error rates. There were significant improvements in the V3-V4 amplicon sequences, which may have a lower resolution than the full-length amplicon sequences. Therefore, integration of MWTC with GTDB as the database could be used in future studies.

In the cases of NCBI RefSeq and RDP, both classification count and fully classified ratio showed a significant increased by MWTC, accompanied by a substantial reduction in error rates. Although the two databases showed lower classification counts by up to two to three times at the genus and species levels compared to other databases when UWTC was used, applying MWTC narrowed the gap in classification counts, improving to a level achieved by the other databases. Furthermore, the error rate was remarkably enhanced, exceeding the performance of the other databases. The NCBI RefSeq used in this analysis contained only 16S rRNA gene sequences annotated at the species level, excluding the genomes of uncultured microbes. Similarly, RDP was derived from genomic data provided by NCBI GenBank, EMBL, and DDBJ [46], and its results were highly consistent with those of NCBI RefSeq. Thus, we confirmed that using a weighted taxonomy classifier is highly effective, especially in those that exclude uncultured genomes, and its impact becomes even more pronounced when applied to species-specific analyses, such as ruminants. Unlike other databases that predominantly contain uncultured microbes, these NCBI RefSeq and RDP enable the assignment of microbial individuals to lower taxonomic levels, thereby being suitable for increasingly detailed studies on microbes and their communities.

## Conclusion

Overall, the use of MWTC improved taxonomic classification resolution in 16S rRNA gene-based microbiome analysis over AWTC, which applies taxonomic weights based on the EMPO database. Notably, the current study confirmed that taxonomic weights could be applied using both shotgun metagenome and amplicon sequence datasets. In particular, the NCBI RefSeq, known for its strict taxonomy and low resolution, showed a significant improvement in taxonomic resolution when MWTC was applied. This enhancement facilitated species-level microbial analysis, making it more effective than other databases that rely on ambiguous annotations. Thus, we recommend using NCBI RefSeq for species-level classification, and its performance will significantly improve when applying MWTC.

## List of abbreviations

EMPO: Earth Microbiome Project Ontology
PERMANOVA: Permutational multivariate analysis of variance
UWTC: Unweighted taxonomy classifier
AWTC: Average weighted taxonomy classifier
MWTC: Manually weighted taxonomy classifier
GG2: most recent Greengenes database (GG2)

## Data availability

All codes and prebuilt classifiers used for data analysis are available at https://github.com/6seok/rumanclass.

## Acknowledgments

Special thanks to Prof. Jakyeom Seo and Dr. Hanbeen Kim for providing the Hanwoo amplicon sequencing dataset used for validation.

## Declarations

### Competing Interests

The authors declare no competing interests.

### Author’s Contributions

R. K. contributed to the conception, methodology, and data curation and wrote the original manuscript. Z. Y. and T. P. reviewed and edited the manuscript. All authors reviewed the final version.

### Author details

^1^ Department of Animal Science and Technology, Chung-Ang University, Anseong-si, Gyeonggi-do 17546, Republic of Korea

^2^ Department of Animal Sciences, The Ohio State University, Columbus, OH 43210, USA

## Notes

https://github.com/6seok/rumanclass/

### Competing Interest Statement

The authors have declared no competing interest.

