## Supplementary Tables S1 to S11 for "Manually weighted taxonomy classifiers improve species-specific rumen microbiome analysis compared to unweighted or average weighted taxonomy classifiers"

**Supplementary Table**

| **Table S1.** Average values and statistical analysis results of alpha diversity for the *in vivo* dataset at genus level. | | | | | | | | | |
| --- | --- | --- | --- | --- | --- | --- | --- | --- | --- |
|  | **(A)** Based on the full-length amplicon sequences | | | |  | **(B)** Based on the V3-V4 amplicon sequences | | | |
| GG2 2022.10 | | | | | | | | | |
|  | UWTC | AWTC | MWTC | *P*-value |  | UWTC | AWTC | MWTC | *P*-value |
| Evenness | 0.549 | 0.549 | 0.549 | 0.8263 |  | 0.556 | 0.557 | 0.555 | 0.302 |
| Observed features | 42.106^a^ | 41.681^b^ | 42.106^a^ | 0.0489 |  | 188.300^a^ | 177.133^b^ | 170.733^c^ | <0.0001 |
| Shannon index | 2.937 | 2.932 | 2.937 | 0.2219 |  | 4.200^a^ | 4.161^b^ | 4.118^c^ | <0.0001 |
| Simpson index | 0.698^a^ | 0.697^b^ | 0.697^b^ | 0.0053 |  | 0.803^b^ | 0.807^a^ | 0.799^c^ | <0.0001 |
| Inverse Simpson index | 3.815^a^ | 3.814^ab^ | 3.800^b^ | 0.0335 |  | 5.972^b^ | 6.171^a^ | 5.824^c^ | <0.0001 |
| GG2 2024.09 | | | | | | | | | |
|  | UWTC | AWTC | MWTC | *P*-value |  | UWTC | AWTC | MWTC | *P*-value |
| Evenness | 0.549 | 0.550 | 0.549 | 0.4086 |  | 0.554 | 0.555 | 0.555 | 0.8678 |
| Observed features | 42.234^a^ | 41.596^ab^ | 41.319^b^ | 0.0009 |  | 191.467^a^ | 177.900^b^ | 171.767^c^ | <0.0001 |
| Shannon index | 2.939^a^ | 2.933^ab^ | 2.924^b^ | 0.0018 |  | 4.205^a^ | 4.151^b^ | 4.120^c^ | <0.0001 |
| Simpson index | 0.698^a^ | 0.697^b^ | 0.697^b^ | 0.0122 |  | 0.803^b^ | 0.806^a^ | 0.800^c^ | <0.0001 |
| Inverse Simpson index | 3.816^a^ | 3.816^ab^ | 3.800^b^ | 0.0452 |  | 5.965^ab^ | 6.122^a^ | 5.853^b^ | 0.0007 |
| GTDB 220.0 | | | | | | | | | |
|  | UWTC | AWTC | MWTC | *P*-value |  | UWTC | AWTC | MWTC | *P*-value |
| Evenness | 0.559^a^ | 0.551^b^ | 0.552^b^ | <0.0001 |  | 0.549^a^ | 0.546^b^ | 0.551^a^ | <0.0001 |
| Observed features | 41.128^a^ | 39.596^c^ | 40.255^b^ | <0.0001 |  | 180.867^a^ | 165.800^c^ | 176.233^b^ | <0.0001 |
| Shannon index | 2.973^a^ | 2.896^b^ | 2.916^b^ | <0.0001 |  | 4.122^a^ | 4.029^b^ | 4.112^a^ | <0.0001 |
| Simpson index | 0.704^a^ | 0.697^b^ | 0.699^b^ | <0.0001 |  | 0.801^a^ | 0.798^b^ | 0.799^b^ | <0.0001 |
| Inverse Simpson index | 3.976^a^ | 3.799^b^ | 3.846^b^ | <0.0001 |  | 5.844^a^ | 5.759^b^ | 5.771^b^ | <0.0001 |
| NCBI 2024.10 | | | | | | | | | |
|  | UWTC | AWTC | MWTC | *P*-value |  | UWTC | AWTC | MWTC | *P*-value |
| Evenness | 0.617^b^ | 0.669^a^ | 0.576^c^ | <0.0001 |  | 0.617^b^ | 0.634^a^ | 0.628^a^ | <0.0001 |
| Observed features | 28.149^b^ | 28.809^b^ | 30.043^a^ | <0.0001 |  | 99.367^b^ | 100.333^b^ | 107.167^a^ | <0.0001 |
| Shannon index | 2.953^b^ | 3.217^a^ | 2.810^c^ | <0.0001 |  | 4.096^b^ | 4.211^a^ | 4.232^a^ | <0.0001 |
| Simpson index | 0.765^b^ | 0.819^a^ | 0.719^c^ | <0.0001 |  | 0.880^b^ | 0.896^a^ | 0.880^b^ | <0.0001 |
| Inverse Simpson index | 4.597^b^ | 5.978^a^ | 3.873^c^ | <0.0001 |  | 9.153^b^ | 10.583^a^ | 9.452^b^ | <0.0001 |
| RDP 2.14 | | | | | | | | | |
|  | UWTC | AWTC | MWTC | *P*-value |  | UWTC | AWTC | MWTC | *P*-value |
| Evenness | 0.613^b^ | 0.663^a^ | 0.580^c^ | <0.0001 |  | 0.603^b^ | 0.631^a^ | 0.629^a^ | <0.0001 |
| Observed features | 25.489^c^ | 27.596^b^ | 29.298^a^ | <0.0001 |  | 93.167^c^ | 95.733^b^ | 101.433^a^ | <0.0001 |
| Shannon index | 2.843^b^ | 3.149^a^ | 2.813^b^ | <0.0001 |  | 3.941^c^ | 4.149^b^ | 4.193^a^ | <0.0001 |
| Simpson index | 0.763^b^ | 0.812^a^ | 0.721^c^ | <0.0001 |  | 0.876^b^ | 0.897^a^ | 0.879^b^ | <0.0001 |
| Inverse Simpson index | 4.472^b^ | 5.746^a^ | 3.903^c^ | <0.0001 |  | 8.651^c^ | 10.496^a^ | 9.334^b^ | <0.0001 |
| SILVA 138.1 | | | | | | | | | |
|  | UWTC | AWTC | MWTC | *P*-value |  | UWTC | AWTC | MWTC | *P*-value |
| Evenness | 0.566^ab^ | 0.568^a^ | 0.563^b^ | 0.0061 |  | 0.609^a^ | 0.591^b^ | 0.594^b^ | <0.0001 |
| Observed features | 37.043^b^ | 37.383^a^ | 36.915^b^ | 0.0011 |  | 150.233^a^ | 131.167^c^ | 144.133^b^ | <0.0001 |
| Shannon index | 2.925^ab^ | 2.943^a^ | 2.908^b^ | 0.0001 |  | 4.405^a^ | 4.160^c^ | 4.262^b^ | <0.0001 |
| Simpson index | 0.710^ab^ | 0.713^a^ | 0.707^b^ | 0.0005 |  | 0.850^a^ | 0.830^b^ | 0.829^b^ | <0.0001 |
| Inverse Simpson index | 3.914^ab^ | 3.934^a^ | 3.871^b^ | 0.0011 |  | 8.101^a^ | 7.115^b^ | 6.822^b^ | <0.0001 |
| SILVA 138.2 | | | | | | | | | |
|  | UWTC | AWTC | MWTC | *P*-value |  | UWTC | AWTC | MWTC | *P*-value |
| Evenness | 0.566^b^ | 0.576^a^ | 0.563^b^ | <0.0001 |  | 0.609^b^ | 0.620^a^ | 0.595^c^ | <0.0001 |
| Observed features | 37.064^a^ | 36.481^b^ | 36.872^a^ | <0.0001 |  | 153.000^a^ | 134.000^c^ | 146.800^b^ | <0.0001 |
| Shannon index | 2.925^b^ | 2.971^a^ | 2.908^b^ | <0.0001 |  | 4.423^a^ | 4.380^b^ | 4.286^c^ | <0.0001 |
| Simpson index | 0.710^b^ | 0.715^a^ | 0.707^b^ | <0.0001 |  | 0.848^b^ | 0.879^a^ | 0.830^c^ | <0.0001 |
| Inverse Simpson index | 3.909^b^ | 4.110^a^ | 3.870^b^ | <0.0001 |  | 8.032^b^ | 9.533^a^ | 6.895^c^ | <0.0001 |
| UWTC, unweighted taxonomy classifier; AWTC, average weighted taxonomy classifier; MWTC, manually weighted taxonomy classifier | | | | | | | | | |
| **Table S2.** Average values and statistical analysis results of alpha diversity for the *in vivo* dataset at species level. | | | | | | | | | |
|  | **(A)** Based on the full-length amplicon sequences | | | |  | **(B)** Based on the V3-V4 amplicon sequences | | | |
| GG2 2022.10 | | | | | | | | | |
|  | UWTC | AWTC | MWTC | *P*-value |  | UWTC | AWTC | MWTC | *P*-value |
| Evenness | 0.700 | 0.696 | 0.695 | 0.1102 |  | 0.638 | 0.638 | 0.641 | 0.0986 |
| Observed features | 57.255^a^ | 56.277^b^ | 56.043^b^ | 0.0006 |  | 277.500^a^ | 259.833^b^ | 250.000^c^ | <0.0001 |
| Shannon index | 4.048^a^ | 4.008^b^ | 3.993^b^ | 0.0025 |  | 5.173^a^ | 5.115^b^ | 5.102^b^ | <0.0001 |
| Simpson index | 0.872 | 0.866 | 0.865 | 0.0779 |  | 0.907^b^ | 0.909^a^ | 0.907^b^ | 0.0466 |
| Inverse Simpson index | 8.995^a^ | 8.677^b^ | 8.690^b^ | 0.0479 |  | 13.089 | 13.044 | 12.806 | 0.1799 |
| GG2 2024.09 | | | | | | | | | |
|  | UWTC | AWTC | MWTC | *P*-value |  | UWTC | AWTC | MWTC | *P*-value |
| Evenness | 0.667^b^ | 0.694^a^ | 0.673^b^ | <0.0001 |  | 0.612^b^ | 0.614^b^ | 0.620^a^ | 0.0042 |
| Observed features | 57.362^a^ | 56.638^ab^ | 55.915^b^ | <0.0001 |  | 282.067^a^ | 259.633^b^ | 250.733^c^ | <0.0001 |
| Shannon index | 3.858^b^ | 3.999^a^ | 3.867^b^ | <0.0001 |  | 4.983^a^ | 4.924^b^ | 4.944^ab^ | 0.0174 |
| Simpson index | 0.840^c^ | 0.865^a^ | 0.850^b^ | <0.0001 |  | 0.866^b^ | 0.872^a^ | 0.871^ab^ | 0.038 |
| Inverse Simpson index | 7.491^b^ | 8.663^a^ | 7.722^b^ | <0.0001 |  | 9.255 | 9.626 | 9.500 | 0.0695 |
| GTDB 220.0 | | | | | | | | | |
|  | UWTC | AWTC | MWTC | *P*-value |  | UWTC | AWTC | MWTC | *P*-value |
| Evenness | 0.660^c^ | 0.668^b^ | 0.680^a^ | <0.0001 |  | 0.621^a^ | 0.606^b^ | 0.630^a^ | <0.0001 |
| Observed features | 54.660 | 53.787 | 53.979 | 0.0526 |  | 241.600^a^ | 224.000^b^ | 243.300^a^ | <0.0001 |
| Shannon index | 3.774^b^ | 3.805^b^ | 3.876^a^ | <0.0001 |  | 4.912^a^ | 4.729^b^ | 4.995^a^ | <0.0001 |
| Simpson index | 0.824^b^ | 0.841^a^ | 0.850^a^ | <0.0001 |  | 0.881^ab^ | 0.864^b^ | 0.888^a^ | 0.0243 |
| Inverse Simpson index | 6.850^c^ | 7.323^b^ | 7.888^a^ | <0.0001 |  | 9.564^ab^ | 8.778^b^ | 10.691^a^ | 0.0032 |
| NCBI 2024.10 | | | | | | | | | |
|  | UWTC | AWTC | MWTC | *P*-value |  | UWTC | AWTC | MWTC | *P*-value |
| Evenness | 0.613^b^ | 0.665^a^ | 0.569^c^ | <0.0001 |  | 0.605^c^ | 0.639^a^ | 0.618^b^ | <0.0001 |
| Observed features | 30.723^b^ | 31.404^b^ | 33.085^a^ | <0.0001 |  | 121.533^c^ | 125.200^b^ | 135.167^a^ | <0.0001 |
| Shannon index | 3.007^b^ | 3.273^a^ | 2.850^c^ | <0.0001 |  | 4.187^c^ | 4.453^a^ | 4.374^b^ | <0.0001 |
| Simpson index | 0.768^b^ | 0.822^a^ | 0.720^c^ | <0.0001 |  | 0.881^b^ | 0.909^a^ | 0.881^b^ | <0.0001 |
| Inverse Simpson index | 4.668^b^ | 6.090^a^ | 3.891^c^ | <0.0001 |  | 9.253^b^ | 12.277^a^ | 9.618^b^ | <0.0001 |
| RDP 2.14 | | | | | | | | | |
|  | UWTC | AWTC | MWTC | *P*-value |  | UWTC | AWTC | MWTC | *P*-value |
| Evenness | 0.609^b^ | 0.658^a^ | 0.574^c^ | <0.0001 |  | 0.590^c^ | 0.637^a^ | 0.619^b^ | <0.0001 |
| Observed features | 27.043^c^ | 29.319^b^ | 31.447^a^ | <0.0001 |  | 111.733^c^ | 117.467^b^ | 127.167^a^ | <0.0001 |
| Shannon index | 2.870^b^ | 3.176^a^ | 2.837^b^ | <0.0001 |  | 4.015^b^ | 4.376^a^ | 4.325^a^ | <0.0001 |
| Simpson index | 0.764^b^ | 0.814^a^ | 0.721^c^ | <0.0001 |  | 0.876^b^ | 0.911^a^ | 0.881^b^ | <0.0001 |
| Inverse Simpson index | 4.507^b^ | 5.787^a^ | 3.918^c^ | <0.0001 |  | 8.702^c^ | 12.152^a^ | 9.505^b^ | <0.0001 |
| SILVA 138.1 | | | | | | | | | |
|  | UWTC | AWTC | MWTC | *P*-value |  | UWTC | AWTC | MWTC | *P*-value |
| Evenness | 0.667^a^ | 0.661^a^ | 0.641^b^ | <0.0001 |  | 0.650^b^ | 0.645^b^ | 0.661^a^ | <0.0001 |
| Observed features | 53.362^a^ | 51.596^b^ | 51.191^b^ | <0.0001 |  | 250.700^a^ | 195.067^c^ | 236.200^b^ | <0.0001 |
| Shannon index | 3.790^a^ | 3.728^b^ | 3.609^c^ | <0.0001 |  | 5.181^a^ | 4.906^b^ | 5.211^a^ | <0.0001 |
| Simpson index | 0.843^a^ | 0.834^a^ | 0.808^b^ | <0.0001 |  | 0.905^b^ | 0.913^a^ | 0.919^a^ | <0.0001 |
| Inverse Simpson index | 7.323^a^ | 6.898^a^ | 6.184^b^ | <0.0001 |  | 12.586^b^ | 13.119^b^ | 14.378^a^ | <0.0001 |
| SILVA 138.2 | | | | | | | | | |
|  | UWTC | AWTC | MWTC | *P*-value |  | UWTC | AWTC | MWTC | *P*-value |
| Evenness | 0.631^b^ | 0.652^a^ | 0.659^a^ | <0.0001 |  | 0.627^b^ | 0.640^a^ | 0.630^ab^ | 0.0156 |
| Observed features | 53.319^a^ | 50.444^b^ | 51.830^b^ | 0.0008 |  | 254.000^a^ | 196.233^c^ | 238.300^b^ | <0.0001 |
| Shannon index | 3.587^b^ | 3.660^ab^ | 3.716^a^ | 0.0004 |  | 5.009^a^ | 4.872^b^ | 4.976^a^ | 0.0006 |
| Simpson index | 0.788^b^ | 0.819^a^ | 0.830^a^ | <0.0001 |  | 0.870^b^ | 0.909^a^ | 0.874^b^ | <0.0001 |
| Inverse Simpson index | 5.844^b^ | 6.697^a^ | 6.813^a^ | <0.0001 |  | 9.924^b^ | 12.861^a^ | 9.259^b^ | <0.0001 |
| UWTC, unweighted taxonomy classifier; AWTC, average weighted taxonomy classifier; MWTC, manually weighted taxonomy classifier | | | | | | | | | |

| **Table S3.** Multiple comparison results for each factor based on differences in microbial composition in the *in vivo* study for the full-length amplicon sequences (A) and the V3-V4 amplicon sequences (B). The Bray-Curtis dissimilarity method was used, and all comparisons were adjusted using the Benjamini-Hochberg method. Pink indicates results at the genus level, and blue indicates results at the species level. | | | | | | | | |
| --- | --- | --- | --- | --- | --- | --- | --- | --- |
| **A** | | | |  | **B** | | | |
| GG2 2022.10 | | | |  | GG2 2022.10 | | | |
|  | UWTC | AWTC | MWTC |  |  | UWTC | AWTC | MWTC |
| UWTC |  | **0.002** | 0.125 |  | UWTC |  | **0.002** | **0.003** |
| AWTC | 0.814 |  | **0.002** |  | AWTC | **0.032** |  | **0.002** |
| MWTC | 0.783 | 0.546 |  |  | MWTC | **0.032** | **0.009** |  |
| GG2 2024.09 | | | |  | GG2 2024.09 | | | |
|  | UWTC | AWTC | MWTC |  |  | UWTC | AWTC | MWTC |
| UWTC |  | **0.002** | **0.009** |  | UWTC |  | **0.002** | **0.002** |
| AWTC | 0.840 |  | **0.002** |  | AWTC | **0.046** |  | **0.002** |
| MWTC | 0.788 | 0.471 |  |  | MWTC | **0.046** | **0.006** |  |
| GTDB 220.0 | | | |  | GTDB 220.0 | | | |
|  | UWTC | AWTC | MWTC |  |  | UWTC | AWTC | MWTC |
| UWTC |  | **0.001** | **0.001** |  | UWTC |  | **0.001** | **0.001** |
| AWTC | 0.762 |  | **0.001** |  | AWTC | **0.009** |  | **0.001** |
| MWTC | 0.845 | 0.844 |  |  | MWTC | **0.009** | **0.006** |  |
| NCBI 2024.10 | | | |  | NCBI 2024.10 | | | |
|  | UWTC | AWTC | MWTC |  |  | UWTC | AWTC | MWTC |
| UWTC |  | **0.001** | **0.001** |  | UWTC |  | **0.001** | **0.001** |
| AWTC | **0.001** |  | **0.001** |  | AWTC | **0.001** |  | **0.001** |
| MWTC | **0.001** | **0.001** |  |  | MWTC | **0.001** | **0.001** |  |
| RDP 2.14 | | | |  | RDP 2.14 | | | |
|  | UWTC | AWTC | MWTC |  |  | UWTC | AWTC | MWTC |
| UWTC |  | **0.001** | **0.001** |  | UWTC |  | **0.001** | **0.001** |
| AWTC | **0.001** |  | **0.001** |  | AWTC | **0.001** |  | **0.001** |
| MWTC | **0.001** | **0.001** |  |  | MWTC | **0.001** | **0.001** |  |
| SILVA 138.1 | | | |  | SILVA 138.1 | | | |
|  | UWTC | AWTC | MWTC |  |  | UWTC | AWTC | MWTC |
| UWTC |  | **0.001** | **0.001** |  | UWTC |  | **0.001** | **0.001** |
| AWTC | 1 |  | **0.001** |  | AWTC | **0.002** |  | **0.001** |
| MWTC | 1 | 1 |  |  | MWTC | **0.022** | **0.002** |  |
| SILVA 138.2 | | | |  | SILVA 138.2 | | | |
|  | UWTC | AWTC | MWTC |  |  | UWTC | AWTC | MWTC |
| UWTC |  | **0.002** | **0.002** |  | UWTC |  | **0.001** | **0.001** |
| AWTC | 1 |  | **0.002** |  | AWTC | **0.002** |  | **0.001** |
| MWTC | 1 | 1 |  |  | MWTC | **0.039** | **0.002** |  |
| UWTC, unweighted taxonomy classifier; AWTC, average weighted taxonomy classifier; MWTC, manually weighted taxonomy classifier | | | | | | | | |

| **Table S4.** Average values and statistical analysis results of classification counts for the *in vitro* dataset. The percentage differences were calculated to indicate how much AWTC and MWTC differed from UWTC. | | | | | | | | | | | | | | | | | | | | | | |
| --- | --- | --- | --- | --- | --- | --- | --- | --- | --- | --- | --- | --- | --- | --- | --- | --- | --- | --- | --- | --- | --- | --- |
| (1) Based on the full-length amplicon sequences | | | | | | | | | | | | | | | | | | | | | | |
|  | Measurements (%) | | | | | | | | | | | | | | |  |  | |  | |  |  |
| Taxonomy | Phylum | | |  | Family | | |  | Genus | | |  | Species | | |  | *P*-value | | | | | |
| Classifier ^1)^ | UWTC | AWTC | MWTC |  | UWTC | AWTC | MWTC |  | UWTC | AWTC | MWTC |  | UWTC | AWTC | MWTC |  | P ^2)^ | F | | G | | S |
| GG2 2022.10 | 147.30^a^ | 146.83^b^ | 146.43^c^ |  | 145.66^a^ | 145.04^b^ | 145.06^b^ |  | 133.74^a^ | 130.74^b^ | 134.15^a^ |  | 94.51^b^ | 93.74^b^ | 99.38^a^ |  | < 0.0001 | < 0.0001 | | < 0.0001 | | < 0.0001 |
|  |  | -0.32 | -0.59 |  |  | -0.42 | -0.41 |  |  | -2.24 | 0.30 |  |  | -0.81 | 5.16 |  |  |  | |  | |  |
| GG2 2024.09 | 147.17^a^ | 146.87^b^ | 146.40^c^ |  | 145.11^a^ | 144.74^b^ | 144.77^b^ |  | 133.74^a^ | 130.55^b^ | 133.89^a^ |  | 83.62^c^ | 86.00^b^ | 90.13^a^ |  | < 0.0001 | 0.0209 | | < 0.0001 | | < 0.0001 |
|  |  | -0.20 | -0.52 |  |  | -0.25 | -0.23 |  |  | -2.39 | 0.11 |  |  | 2.85 | 7.79 |  |  |  | |  | |  |
| GTDB 220.0 | 146.81 | 147.02 | 147.04 |  | 144.04^b^ | 144.83^b^ | 145.94^a^ |  | 125.85^c^ | 127.96^b^ | 130.72^a^ |  | 58.47^b^ | 74.19^a^ | 73.06^a^ |  | 0.1987 | < 0.0001 | | < 0.0001 | | < 0.0001 |
|  |  | 0.14 | 0.16 |  |  | 0.55 | 1.31 |  |  | 1.67 | 3.87 |  |  | 26.89 | 24.96 |  |  |  | |  | |  |
| NCBI 2024.10 | 144.06^c^ | 144.72^b^ | 145.51^a^ |  | 98.87^b^ | 105.60^a^ | 105.74^a^ |  | 68.00^c^ | 73.51^b^ | 85.32^a^ |  | 58.74^c^ | 65.15^b^ | 76.55^a^ |  | < 0.0001 | < 0.0001 | | < 0.0001 | | < 0.0001 |
|  |  | 0.46 | 1.00 |  |  | 6.80 | 6.95 |  |  | 8.10 | 25.47 |  |  | 10.90 | 30.32 |  |  |  | |  | |  |
| RDP  2.14 | 145.85^b^ | 145.98^b^ | 146.45^a^ |  | 100.32^c^ | 108.40^b^ | 114.19^a^ |  | 60.91^c^ | 71.74^b^ | 83.49^a^ |  | 55.45^c^ | 63.53^b^ | 75.40^a^ |  | < 0.0001 | < 0.0001 | | < 0.0001 | | < 0.0001 |
|  |  | 0.09 | 0.41 |  |  | 8.06 | 13.83 |  |  | 17.78 | 37.06 |  |  | 14.58 | 35.99 |  |  |  | |  | |  |
| SILVA 138.1 | 147.49^b^ | 147.49^b^ | 147.66^a^ |  | 147.43^a^ | 146.94^b^ | 147.53^a^ |  | 146.06^a^ | 143.83^b^ | 146.28^a^ |  | 95.02^c^ | 83.17^b^ | 110.98^a^ |  | 0.0011 | < 0.0001 | | < 0.0001 | | < 0.0001 |
|  |  | 0.00 | 0.12 |  |  | -0.33 | 0.07 |  |  | -1.53 | 0.15 |  |  | -12.47 | 16.79 |  |  |  | |  | |  |
| SILVA 138.2 | 147.49^b^ | 143.00^b^ | 147.66^a^ |  | 147.43^a^ | 142.41^b^ | 147.57^a^ |  | 145.91^a^ | 138.44^b^ | 146.40^a^ |  | 72.72^c^ | 82.19^b^ | 89.77^a^ |  | 0.0048 | < 0.0001 | | < 0.0001 | | < 0.0001 |
|  |  | -3.04 | 0.12 |  |  | -3.40 | 0.10 |  |  | -5.12 | 0.34 |  |  | 13.01 | 23.43 |  |  |  | |  | |  |
| **(**2) Based on the V3-V4 amplicon sequences | | | | | | | | | | | | | | | | | | | | | | |
|  | Measurements (%) | | | | | | | | | | | | | | |  |  | |  | |  |  |
| Taxonomy | Phylum | | |  | Family | | |  | Genus | | |  | Species | | |  | *P*-value | | | | | |
| Classifier | UWTC | AWTC | MWTC |  | UWTC | AWTC | MWTC |  | UWTC | AWTC | MWTC |  | UWTC | AWTC | MWTC |  | P | | F | | G | S |
| GG2 2022.10 | 890.50^a^ | 887.13^a^ | 850.63^b^ |  | 861.27^a^ | 858.30^a^ | 838.93^b^ |  | 698.97^b^ | 680.40^c^ | 711.47^a^ |  | 510.83^b^ | 513.23^b^ | 548.97^a^ |  | < 0.0001 | | < 0.0001 | | < 0.0001 | < 0.0001 |
|  |  | -0.38 | -4.48 |  |  | -0.34 | -2.59 |  |  | -2.66 | 1.79 |  |  | 0.47 | 7.46 |  |  | |  | |  |  |
| GG2 2024.09 | 889.77^a^ | 877.40^b^ | 846.43^c^ |  | 859.27^a^ | 851.83^b^ | 835.47^c^ |  | 699.63^a^ | 674.10^b^ | 703.60^a^ |  | 495.63^b^ | 483.63^c^ | 525.73^a^ |  | < 0.0001 | | < 0.0001 | | < 0.0001 | < 0.0001 |
|  |  | -1.39 | -4.87 |  |  | -0.87 | -2.77 |  |  | -3.65 | 0.57 |  |  | -2.42 | 6.07 |  |  | |  | |  |  |
| GTDB 220.0 | 878.17^c^ | 883.80^b^ | 890.00^a^ |  | 822.60^c^ | 827.77^b^ | 853.17^a^ |  | 606.93^b^ | 594.23^c^ | 671.17^a^ |  | 294.80^b^ | 302.80^b^ | 406.30^a^ |  | < 0.0001 | | < 0.0001 | | < 0.0001 | < 0.0001 |
|  |  | 0.64 | 1.35 |  |  | 0.63 | 3.72 |  |  | -2.09 | 10.58 |  |  | 2.71 | 37.82 |  |  | |  | |  |  |
| NCBI 2024.10 | 865.17^c^ | 888.33^a^ | 881.07^b^ |  | 582.40^c^ | 603.00^b^ | 651.40^a^ |  | 292.30^c^ | 397.80^b^ | 410.87^a^ |  | 181.60^c^ | 288.93^b^ | 327.97^a^ |  | < 0.0001 | | < 0.0001 | | < 0.0001 | < 0.0001 |
|  |  | 2.68 | 1.84 |  |  | 3.54 | 11.85 |  |  | 36.09 | 40.56 |  |  | 59.10 | 80.60 |  |  | |  | |  |  |
| RDP  2.14 | 876.93^c^ | 889.27^b^ | 898.90^a^ |  | 621.70^c^ | 652.07^b^ | 716.47^a^ |  | 254.77^c^ | 349.33^b^ | 387.80^a^ |  | 178.73^c^ | 248.07^b^ | 307.83^a^ |  | < 0.0001 | | < 0.0001 | | < 0.0001 | < 0.0001 |
|  |  | 1.41 | 2.50 |  |  | 4.88 | 15.24 |  |  | 37.12 | 52.22 |  |  | 38.79 | 72.23 |  |  | |  | |  |  |
| SILVA 138.1 | 913.70^a^ | 913.40^a^ | 906.37^b^ |  | 883.67^b^ | 882.43^b^ | 887.03^a^ |  | 769.07^b^ | 713.07^c^ | 813.83^a^ |  | 337.57^b^ | 518.63^a^ | 511.87^a^ |  | < 0.0001 | | 0.0011 | | < 0.0001 | < 0.0001 |
|  |  | -0.03 | -0.80 |  |  | -0.14 | 0.38 |  |  | -7.28 | 5.82 |  |  | 53.64 | 51.63 |  |  | |  | |  |  |
| SILVA 138.2 | 913.77^a^ | 913.87^a^ | 906.33^b^ |  | 883.80^a^ | 880.63^b^ | 886.40^a^ |  | 774.07^b^ | 662.97^c^ | 813.63^a^ |  | 335.57^b^ | 485.90^a^ | 473.53^a^ |  | < 0.0001 | | < 0.0001 | | < 0.0001 | < 0.0001 |
|  |  | 0.01 | -0.81 |  |  | -0.36 | 0.29 |  |  | -14.35 | 5.11 |  |  | 44.80 | 41.11 |  |  | |  | |  |  |
| ^a-c^ Different superscripts in a row indicate statistically significant differences (*p* < 0.05).  ^1)^ UWTC, unweighted taxonomy classifier; AWTC, average weighted taxonomy classifier; MWTC, manually weighted taxonomy classifier  ^2)^ P, phylum; F, family; G, genus; S, species | | | | | | | | | | | | | | | | | | | | | | |

| **Table S5.** Average values and statistical analysis results of fully-classified ratios for the *in vivo* dataset. The percentage difference was calculated to indicate how much AWTC and MWTC differed from UWTC. | | | | | | | | | | |
| --- | --- | --- | --- | --- | --- | --- | --- | --- | --- | --- |
| (1) Based on the full-length amplicon sequences | | | | | | | | | | |
|  | Measurements (%) | | | | | | |  |  |  |
| Taxonomy level | Genus | | |  | Species | | |  | *P*-value | |
| Classifier ^1)^ | UWTC | AWTC | MWTC |  | UWTC | AWTC | MWTC |  | Genus | Species |
| GG2 2022.10 | 94.49^a^ | 92.99^b^ | 93.86^a^ |  | 70.09 | 67.65 | 72.13 |  | < 0.0001 | 0.0682 |
|  |  | -1.59 | -0.67 |  |  | -3.49 | 2.91 |  |  |  |
| GG2 2024.09 | 94.57^a^ | 92.95^b^ | 93.86^a^ |  | 55.21^b^ | 60.22^a^ | 61.26^a^ |  | < 0.0001 | < 0.0001 |
|  |  | -1.71 | -0.74 |  |  | 9.08 | 10.94 |  |  |  |
| GTDB 220.0 | 91.26^c^ | 92.88^b^ | 93.92^a^ |  | 38.09^c^ | 52.61^a^ | 49.45^b^ |  | < 0.0001 | < 0.0001 |
|  |  | 1.77 | 2.91 |  |  | 38.13 | 29.82 |  |  |  |
| NCBI 2024.10 | 55.18^c^ | 58.95^b^ | 68.23^a^ |  | 50.48^c^ | 54.36^b^ | 63.63^a^ |  | < 0.0001 | < 0.0001 |
|  |  | 6.85 | 23.66 |  |  | 7.69 | 26.06 |  |  |  |
| RDP 2.14 | 49.52^c^ | 57.93^b^ | 67.01^a^ |  | 47.09^c^ | 53.89^b^ | 62.91^a^ |  | < 0.0001 | < 0.0001 |
|  |  | 16.96 | 35.31 |  |  | 14.43 | 33.58 |  |  |  |
| SILVA 138.1 | 99.42^a^ | 98.42^b^ | 99.58^a^ |  | 65.04^b^ | 54.15^c^ | 78.55^a^ |  | < 0.0001 | < 0.0001 |
|  |  | -1.00 | 0.16 |  |  | -16.74 | 20.77 |  |  |  |
| SILVA 138.2 | 99.26^a^ | 97.89^b^ | 99.54^a^ |  | 45.05^c^ | 52.72^b^ | 60.38^a^ |  | < 0.0001 | < 0.0001 |
|  |  | -1.38 | 0.28 |  |  | 17.04 | 34.03 |  |  |  |
| (2) Based on the V3-V4 amplicon sequences | | | | | | | | | | |
|  | Measurements (%) | | | | | | |  |  |  |
| Taxonomy level | Genus | | |  | Species | | |  | *P*-value | |
| Classifier ^1)^ | UWTC | AWTC | MWTC |  | UWTC | AWTC | MWTC |  | Genus | Species |
| GG2 2022.10 | 86.61^b^ | 82.92^c^ | 88.48^a^ |  | 40.19^c^ | 51.38^b^ | 55.48^a^ |  | < 0.0001 | < 0.0001 |
|  |  | -4.26 | 2.16 |  |  | 27.85 | 38.05 |  |  |  |
| GG2 2024.09 | 86.84^b^ | 83.05^c^ | 88.14^a^ |  | 40.19^b^ | 37.45^c^ | 42.33^a^ |  | < 0.0001 | < 0.0001 |
|  |  | -4.36 | 1.50 |  |  | -6.81 | 5.33 |  |  |  |
| GTDB 220.0 | 81.48^b^ | 80.81^c^ | 85.25^a^ |  | 33.83^b^ | 34.75^b^ | 43.93^a^ |  | < 0.0001 | < 0.0001 |
|  |  | -0.83 | 4.62 |  |  | 2.74 | 29.87 |  |  |  |
| NCBI 2024.10 | 44.51^b^ | 59.07^a^ | 58.92^a^ |  | 35.42^c^ | 47.66^b^ | 53.03^a^ |  | < 0.0001 | < 0.0001 |
|  |  | 32.72 | 32.37 |  |  | 34.54 | 49.71 |  |  |  |
| RDP 2.14 | 40.19^c^ | 50.59^b^ | 56.84^a^ |  | 33.82^c^ | 39.69^b^ | 49.81^a^ |  | < 0.0001 | < 0.0001 |
|  |  | 25.88 | 41.42 |  |  | 17.35 | 47.27 |  |  |  |
| SILVA 138.1 | 89.01^b^ | 86.45^c^ | 93.99^a^ |  | 37.27^c^ | 55.87^b^ | 59.70^a^ |  | < 0.0001 | < 0.0001 |
|  |  | -2.87 | 5.59 |  |  | 49.91 | 60.19 |  |  |  |
| SILVA 138.2 | 90.11^b^ | 76.28^c^ | 94.50^a^ |  | 30.71^c^ | 42.60^b^ | 46.59^a^ |  | < 0.0001 | < 0.0001 |
|  |  | -15.35 | 4.88 |  |  | 38.69 | 51.69 |  |  |  |
| ^a-c^ Different superscripts in a row indicate statistically significant differences (*p* < 0.05).  ^1)^ UWTC, unweighted taxonomy classifier; AWTC, average weighted taxonomy classifier; MWTC, manually weighted taxonomy classifier | | | | | | | | | | |

| **Table S6.** Average values and statistical analysis results of alpha diversity for the *in vitro* dataset at genus level. | | | | | | | | | |
| --- | --- | --- | --- | --- | --- | --- | --- | --- | --- |
|  | **(A)** Based on the full-length amplicon sequences | | | |  | **(B)** Based on the V3-V4 amplicon sequences | | | |
| GG2 2022.10 | | | | | | | | | |
|  | UWTC | AWTC | MWTC | *P*-value |  | UWTC | AWTC | MWTC | *P*-value |
| Evenness | 0.780 | 0.778 | 0.780 | 0.8005 |  | 0.797^a^ | 0.722^b^ | 0.711^c^ | <0.0001 |
| Observed features | 131.400^a^ | 126.000^b^ | 127.800^ab^ | 0.0185 |  | 379.250^a^ | 236.750^b^ | 228.500^b^ | <0.0001 |
| Shannon index | 5.488 | 5.426 | 5.458 | 0.1245 |  | 6.821^a^ | 5.692^b^ | 5.566^c^ | <0.0001 |
| Simpson index | 0.958 | 0.957 | 0.958 | 0.7747 |  | 0.955^a^ | 0.955^a^ | 0.950^b^ | <0.0001 |
| Inverse Simpson index | 23.898 | 23.445 | 23.663 | 0.7631 |  | 22.393^a^ | 22.226^a^ | 20.098^b^ | 0.0002 |
| GG2 2024.09 | | | | | | | | | |
|  | UWTC | AWTC | MWTC | *P*-value |  | UWTC | AWTC | MWTC | *P*-value |
| Evenness | 0.777 | 0.781 | 0.780 | 0.741 |  | 0.727^a^ | 0.723^b^ | 0.710^c^ | <0.0001 |
| Observed features | 135.600^a^ | 128.400^b^ | 129.600^b^ | 0.0007 |  | 261.750^a^ | 235.750^b^ | 229.000^c^ | <0.0001 |
| Shannon index | 5.507 | 5.467 | 5.477 | 0.3256 |  | 5.835^a^ | 5.697^b^ | 5.567^c^ | <0.0001 |
| Simpson index | 0.958 | 0.958 | 0.958 | 0.9676 |  | 0.955^a^ | 0.954^a^ | 0.950^b^ | <0.0001 |
| Inverse Simpson index | 23.925 | 23.819 | 23.762 | 0.965 |  | 22.395^a^ | 22.061^a^ | 19.927^b^ | <0.0001 |
| GTDB 220.0 | | | | | | | | | |
|  | UWTC | AWTC | MWTC | *P*-value |  | UWTC | AWTC | MWTC | *P*-value |
| Evenness | 0.803^a^ | 0.788^b^ | 0.783^b^ | 0.0008 |  | 0.706^c^ | 0.711^b^ | 0.715^a^ | 0.0013 |
| Observed features | 125.000 | 123.200 | 124.800 | 0.792 |  | 248.250^a^ | 230.250^b^ | 242.750^a^ | 0.0104 |
| Shannon index | 5.593^a^ | 5.470^b^ | 5.453^b^ | 0.0018 |  | 5.614^b^ | 5.575^c^ | 5.662^a^ | <0.0001 |
| Simpson index | 0.965^a^ | 0.960^b^ | 0.959^b^ | <0.0001 |  | 0.951^a^ | 0.950^b^ | 0.951^a^ | 0.0126 |
| Inverse Simpson index | 28.770^a^ | 24.989^b^ | 24.140^b^ | <0.0001 |  | 20.686^a^ | 20.294^b^ | 20.692^a^ | 0.0207 |
| NCBI 2024.10 | | | | | | | | | |
|  | UWTC | AWTC | MWTC | *P*-value |  | UWTC | AWTC | MWTC | *P*-value |
| Evenness | 0.772^a^ | 0.752^b^ | 0.764^ab^ | 0.0491 |  | 0.723^c^ | 0.737^b^ | 0.757^a^ | <0.0001 |
| Observed features | 77.200^b^ | 78.200^b^ | 83.600^a^ | 0.0174 |  | 117.500^b^ | 118.250^ab^ | 121.750^a^ | 0.0379 |
| Shannon index | 4.840^a^ | 4.730^b^ | 4.878^a^ | 0.00037 |  | 4.972^c^ | 5.074^b^ | 5.247^a^ | <0.0001 |
| Simpson index | 0.945^a^ | 0.929^b^ | 0.933^b^ | <0.0001 |  | 0.948^c^ | 0.955^b^ | 0.961^a^ | <0.0001 |
| Inverse Simpson index | 18.145^a^ | 14.112^b^ | 14.941^b^ | <0.0001 |  | 19.198^c^ | 22.357^b^ | 25.530^a^ | <0.0001 |
| RDP 2.14 | | | | | | | | | |
|  | UWTC | AWTC | MWTC | *P*-value |  | UWTC | AWTC | MWTC | *P*-value |
| Evenness | 0.730^b^ | 0.726^b^ | 0.758^a^ | 0.0008 |  | 0.711^c^ | 0.722^b^ | 0.762^a^ | <0.0001 |
| Observed features | 69.800^b^ | 73.000^b^ | 77.200^a^ | 0.0013 |  | 110.500^b^ | 117.500^a^ | 114.750^a^ | 0.0014 |
| Shannon index | 4.468^b^ | 4.496^b^ | 4.752^a^ | <0.0001 |  | 4.824^c^ | 4.963^b^ | 5.213^a^ | <0.0001 |
| Simpson index | 0.914^b^ | 0.914^b^ | 0.929^a^ | 0.0003 |  | 0.943^c^ | 0.951^b^ | 0.959^a^ | <0.0001 |
| Inverse Simpson index | 11.676^b^ | 11.658^b^ | 14.113^a^ | 0.0001 |  | 17.693^c^ | 20.248^b^ | 24.474^a^ | <0.0001 |
| SILVA 138.1 | | | | | | | | | |
|  | UWTC | AWTC | MWTC | *P*-value |  | UWTC | AWTC | MWTC | *P*-value |
| Evenness | 0.796 | 0.799 | 0.798 | 0.7892 |  | 0.765^a^ | 0.750^b^ | 0.765^a^ | <0.0001 |
| Observed features | 102.000 | 98.000 | 99.200 | 0.1703 |  | 180.250^a^ | 153.750^c^ | 171.750^b^ | <0.0001 |
| Shannon index | 5.312 | 5.286 | 5.289 | 0.7233 |  | 5.731^a^ | 5.450^c^ | 5.682^b^ | <0.0001 |
| Simpson index | 0.957 | 0.956 | 0.956 | 0.9819 |  | 0.962^a^ | 0.954^c^ | 0.960^b^ | <0.0001 |
| Inverse Simpson index | 23.144 | 23.005 | 23.025 | 0.9808 |  | 26.392^a^ | 21.974^c^ | 25.099^b^ | <0.0001 |
| SILVA 138.2 | | | | | | | | | |
|  | UWTC | AWTC | MWTC | *P*-value |  | UWTC | AWTC | MWTC | *P*-value |
| Evenness | 0.793 | 0.796 | 0.794 | 0.6723 |  | 0.767^a^ | 0.760^b^ | 0.767^a^ | 0.0006 |
| Observed features | 105.400 | 102.600 | 102.600 | 0.362 |  | 184.750^a^ | 153.250^c^ | 174.250^b^ | <0.0001 |
| Shannon index | 5.329 | 5.321 | 5.301 | 0.7702 |  | 5.773^a^ | 5.517^c^ | 5.708^b^ | <0.0001 |
| Simpson index | 0.957 | 0.957 | 0.956 | 0.965 |  | 0.962^a^ | 0.959^c^ | 0.960^b^ | <0.0001 |
| Inverse Simpson index | 23.208 | 23.183 | 23.011 | 0.9638 |  | 26.652^a^ | 24.262^c^ | 25.190^b^ | 0.0003 |
| UWTC, unweighted taxonomy classifier; AWTC, average weighted taxonomy classifier; MWTC, manually weighted taxonomy classifier | | | | | | | | | |

| **Table S7.** AWTC values and statistical analysis results of alpha diversity for the *in vitro* dataset at species level. | | | | | | | | | |
| --- | --- | --- | --- | --- | --- | --- | --- | --- | --- |
|  | **(A)** Based on the full-length amplicon sequences | | | |  | **(B)** Based on the V3-V4 amplicon sequences | | | |
| GG2 2022.10 | | | | | | | | | |
|  | UWTC | AWTC | MWTC | *P*-value |  | UWTC | AWTC | MWTC | *P*-value |
| Evenness | 0.834 | 0.836 | 0.834 | 0.1249 |  | 0.726^b^ | 0.785^a^ | 0.778^a^ | <0.0001 |
| Observed features | 185.600^a^ | 178.000^b^ | 181.600^ab^ | 0.0063 |  | 257.250^c^ | 346.250^a^ | 337.500^b^ | <0.0001 |
| Shannon index | 6.286^a^ | 6.246^b^ | 6.255^b^ | 0.0016 |  | 5.813^c^ | 6.619^a^ | 6.529^b^ | <0.0001 |
| Simpson index | 0.977^b^ | 0.977^a^ | 0.976^c^ | 0.0002 |  | 0.981^a^ | 0.978^b^ | 0.975^c^ | <0.0001 |
| Inverse Simpson index | 42.823^b^ | 43.728^a^ | 42.049^c^ | 0.0001 |  | 52.297^a^ | 45.102^b^ | 39.836^c^ | <0.0001 |
| GG2 2024.09 | | | | | | | | | |
|  | UWTC | AWTC | MWTC | *P*-value |  | UWTC | AWTC | MWTC | *P*-value |
| Evenness | 0.829^b^ | 0.838^a^ | 0.832^b^ | 0.0005 |  | 0.786^a^ | 0.777^b^ | 0.770^c^ | <0.0001 |
| Observed features | 188.800^a^ | 183.000^b^ | 182.400^b^ | 0.0216 |  | 383.000^a^ | 344.750^b^ | 335.750^b^ | <0.0001 |
| Shannon index | 6.267^b^ | 6.297^a^ | 6.251^b^ | 0.0051 |  | 6.739^a^ | 6.551^b^ | 6.456^c^ | <0.0001 |
| Simpson index | 0.976^b^ | 0.978^a^ | 0.976^b^ | <0.0001 |  | 0.977^a^ | 0.974^b^ | 0.971^c^ | <0.0001 |
| Inverse Simpson index | 41.875^b^ | 44.904^a^ | 41.601^b^ | <0.0001 |  | 43.352^a^ | 38.318^b^ | 35.100^c^ | <0.0001 |
| GTDB 220.0 | | | | | | | | | |
|  | UWTC | AWTC | MWTC | *P*-value |  | UWTC | AWTC | MWTC | *P*-value |
| Evenness | 0.833^a^ | 0.828^b^ | 0.818^c^ | <0.0001 |  | 0.765^b^ | 0.759^c^ | 0.777^a^ | <0.0001 |
| Observed features | 171.600 | 172.400 | 174.000 | 0.5356 |  | 327.000^a^ | 308.000^b^ | 328.500^a^ | 0.004 |
| Shannon index | 6.179^b^ | 6.148^b^ | 6.084^a^ | 0.0001 |  | 6.384^b^ | 6.275^c^ | 6.489^a^ | <0.0001 |
| Simpson index | 0.977^a^ | 0.975^b^ | 0.973^c^ | <0.0001 |  | 0.972^b^ | 0.967^c^ | 0.975^a^ | <0.0001 |
| Inverse Simpson index | 44.336^a^ | 39.957^b^ | 37.161^c^ | <0.0001 |  | 36.245^b^ | 30.785^c^ | 40.245^a^ | <0.0001 |
| NCBI 2024.10 | | | | | | | | | |
|  | UWTC | AWTC | MWTC | *P*-value |  | UWTC | AWTC | MWTC | *P*-value |
| Evenness | 0.768^a^ | 0.754^c^ | 0.761^b^ | 0.0008 |  | 0.715^c^ | 0.746^a^ | 0.759^b^ | <0.0001 |
| Observed features | 93.800^b^ | 94.600^b^ | 102.000^a^ | 0.0001 |  | 145.500^b^ | 145.750^b^ | 152.000^a^ | 0.0435 |
| Shannon index | 5.034^b^ | 4.950^c^ | 5.078^a^ | <0.0001 |  | 5.138^c^ | 5.362^b^ | 5.503^a^ | <0.0001 |
| Simpson index | 0.948^a^ | 0.933^c^ | 0.936^b^ | <0.0001 |  | 0.950^c^ | 0.960^b^ | 0.963^a^ | <0.0001 |
| Inverse Simpson index | 19.167^a^ | 14.890^c^ | 15.614^b^ | <0.0001 |  | 19.843^c^ | 25.246^b^ | 27.324^a^ | <0.0001 |
| RDP 2.14 | | | | | | | | | |
|  | UWTC | AWTC | MWTC | *P*-value |  | UWTC | AWTC | MWTC | *P*-value |
| Evenness | 0.719^b^ | 0.725^b^ | 0.751^a^ | <0.0001 |  | 0.704^c^ | 0.728^b^ | 0.758^a^ | <0.0001 |
| Observed features | 85.200^b^ | 89.600^ab^ | 94.000^a^ | 0.0017 |  | 134.000^b^ | 143.000^a^ | 144.750^a^ | 0.0012 |
| Shannon index | 4.609^c^ | 4.699^b^ | 4.919^a^ | <0.0001 |  | 4.976^c^ | 5.209^b^ | 5.439^a^ | <0.0001 |
| Simpson index | 0.916^b^ | 0.917^b^ | 0.931^a^ | <0.0001 |  | 0.945^c^ | 0.955^b^ | 0.961^a^ | <0.0001 |
| Inverse Simpson index | 11.937^b^ | 12.070^b^ | 14.563^a^ | <0.0001 |  | 18.203^c^ | 22.110^b^ | 25.872^a^ | <0.0001 |
| SILVA 138.1 | | | | | | | | | |
|  | UWTC | AWTC | MWTC | *P*-value |  | UWTC | AWTC | MWTC | *P*-value |
| Evenness | 0.819^b^ | 0.832^a^ | 0.827^a^ | 0.0006 |  | 0.794^b^ | 0.769^c^ | 0.810^a^ | <0.0001 |
| Observed features | 175.600^a^ | 160.400^b^ | 157.600^b^ | <0.0001 |  | 326.250^a^ | 248.500^c^ | 291.750^b^ | <0.0001 |
| Shannon index | 6.108^a^ | 6.091^a^ | 6.039^b^ | 0.0013 |  | 6.625^a^ | 6.118^b^ | 6.633^a^ | <0.0001 |
| Simpson index | 0.971^b^ | 0.974^a^ | 0.971^b^ | <0.0001 |  | 0.979^b^ | 0.967^c^ | 0.981^a^ | <0.0001 |
| Inverse Simpson index | 34.044^b^ | 38.670^a^ | 34.133^b^ | <0.0001 |  | 48.259^b^ | 30.803^c^ | 52.145^a^ | <0.0001 |
| SILVA 138.2 | | | | | | | | | |
|  | UWTC | AWTC | MWTC | *P*-value |  | UWTC | AWTC | MWTC | *P*-value |
| Evenness | 0.820^b^ | 0.834^a^ | 0.822^b^ | <0.0001 |  | 0.796^b^ | 0.767^c^ | 0.807^a^ | <0.0001 |
| Observed features | 179.000^a^ | 165.000^b^ | 163.400^b^ | <0.0001 |  | 335.000^a^ | 248.000^c^ | 298.250^b^ | <0.0001 |
| Shannon index | 6.137^a^ | 6.140^a^ | 6.041^b^ | <0.0001 |  | 6.676^a^ | 6.102^c^ | 6.629^b^ | <0.0001 |
| Simpson index | 0.971^b^ | 0.975^a^ | 0.970^b^ | <0.0001 |  | 0.980^a^ | 0.966^b^ | 0.979^a^ | <0.0001 |
| Inverse Simpson index | 34.591^b^ | 39.399^a^ | 33.592^c^ | <0.0001 |  | 49.130^a^ | 29.941^b^ | 47.154^a^ | <0.0001 |
| UWTC, unweighted taxonomy classifier; AWTC, average weighted taxonomy classifier; MWTC, manually weighted taxonomy classifier | | | | | | | | | |

| **Table S8.** Multiple comparison results for each factor based on differences in microbial composition in the *in vitro* study for the full-length amplicon sequences (A) and the V3-V4 amplicon sequences (B). The Bray-Curtis dissimilarity method was used, and all comparisons were adjusted using the Benjamini-Hochberg method. Pink indicates results at the genus level, and blue indicates results at the species level. | | | | | | | | |
| --- | --- | --- | --- | --- | --- | --- | --- | --- |
| **A** | | | |  | **B** | | | |
| GG2-2022.10 | | | |  | GG2-2022.10 | | | |
|  | UWTC | AWTC | MWTC |  |  | UWTC | AWTC | MWTC |
| UWTC |  | **0.009** | **0.009** |  | UWTC |  | **0.032** | **0.032** |
| AWTC | **0.013** |  | **0.009** |  | AWTC | **0.031** |  | **0.032** |
| MWTC | 0.227 | **0.013** |  |  | MWTC | **0.031** | **0.031** |  |
| GG2-2024.09 | | | |  | GG2-2024.09 | | | |
|  | UWTC | AWTC | MWTC |  |  | UWTC | AWTC | MWTC |
| UWTC |  | **0.009** | **0.009** |  | UWTC |  | **0.034** | **0.034** |
| AWTC | **0.015** |  | **0.009** |  | AWTC | **0.03** |  | **0.034** |
| MWTC | 0.212 | **0.015** |  |  | MWTC | **0.03** | **0.03** |  |
| GTDB-220.0 | | | |  | GTDB-220.0 | | | |
|  | UWTC | AWTC | MWTC |  |  | UWTC | AWTC | MWTC |
| UWTC |  | **0.011** | **0.011** |  | UWTC |  | **0.036** | **0.36** |
| AWTC | **0.01** |  | **0.011** |  | AWTC | **0.039** |  | **0.036** |
| MWTC | **0.01** | **0.01** |  |  | MWTC | **0.039** | **0.039** |  |
| NCBI-2024.10 | | | |  | NCBI-2024.10 | | | |
|  | UWTC | AWTC | MWTC |  |  | UWTC | AWTC | MWTC |
| UWTC |  | **0.011** | **0.011** |  | UWTC |  | **0.045** | **0.045** |
| AWTC | **0.014** |  | **0.011** |  | AWTC | **0.03** |  | **0.047** |
| MWTC | **0.014** | **0.014** |  |  | MWTC | **0.03** | **0.03** |  |
| RDP-2.14 | | | |  | RDP-2.14 | | | |
|  | UWTC | AWTC | MWTC |  |  | UWTC | AWTC | MWTC |
| UWTC |  | **0.009** | **0.009** |  | UWTC |  | **0.035** | **0.035** |
| AWTC | **0.011** |  | **0.009** |  | AWTC | **0.04** |  | **0.035** |
| MWTC | **0.011** | **0.011** |  |  | MWTC | **0.04** | **0.04** |  |
| SILVA-138.1 | | | |  | SILVA-138.1 | | | |
|  | UWTC | AWTC | MWTC |  |  | UWTC | AWTC | MWTC |
| UWTC |  | **0.011** | **0.011** |  | UWTC |  | 0.055 | **0.038** |
| AWTC | 0.683 |  | **0.011** |  | AWTC | **0.045** |  | **0.038** |
| MWTC | 0.683 | 0.683 |  |  | MWTC | 0.119 | **0.045** |  |
| SILVA-138.2 | | | |  | SILVA-138.2 | | | |
|  | UWTC | AWTC | MWTC |  |  | UWTC | AWTC | MWTC |
| UWTC |  | **0.01** | **0.01** |  | UWTC |  | **0.036** | **0.036** |
| AWTC | 0.683 |  | **0.01** |  | AWTC | **0.01** |  | **0.036** |
| MWTC | 0.683 | 0.683 |  |  | MWTC | **0.01** | **0.01** |  |
| UWTC, unweighted taxonomy classifier; AWTC, average weighted taxonomy classifier; MWTC, manually weighted taxonomy classifier | | | | | | | | |

| **Table S9.** Average values and statistical analysis results of classification counts for the *in vivo* dataset. The percentage difference was calculated to indicate how much AWTC and MWTC differed from UWTC. | | | | | | | | | | | | | | | | | | | | | | |
| --- | --- | --- | --- | --- | --- | --- | --- | --- | --- | --- | --- | --- | --- | --- | --- | --- | --- | --- | --- | --- | --- | --- |
| (1) Based on full-length amplicon sequences | | | | | | | | | | | | | | | | | | | | | | |
|  | Measurements (%) | | | | | | | | | | | | | | |  |  | |  | |  |  |
| Taxonomy | Phylum | | |  | Family | | |  | Genus | | |  | Species | | |  | *P*-value | | | | | |
| Classifier ^1)^ | UWTC | AWTC | MWTC |  | UWTC | AWTC | MWTC |  | UWTC | AWTC | MWTC |  | UWTC | AWTC | MWTC |  | P ^2)^ | F | | G | | S |
| GG2 2022.10 | 458.20^a^ | 456.00^b^ | 455.00^b^ |  | 453.60^a^ | 452.00^a^ | 450.60^b^ |  | 387.20^a^ | 370.00^b^ | 384.40^a^ |  | 274.60^a^ | 265.00^b^ | 275.80^a^ |  | 0.0004 | 0.0021 | | < 0.0001 | | 0.0001 |
|  |  | -0.48 | -0.70 |  |  | -0.35 | -0.66 |  |  | -4.44 | -0.72 |  |  | -3.50 | 0.44 |  |  |  | |  | |  |
| GG2 2024.09 | 458.20^a^ | 454.40^b^ | 453.60^c^ |  | 453.00^a^ | 449.60^a^ | 449.00^b^ |  | 383.60^a^ | 365.40^c^ | 377.80^b^ |  | 273.00^a^ | 265.00^b^ | 272.60^a^ |  | < 0.0001 | < 0.0001 | | < 0.0001 | | 0.0021 |
|  |  | -0.83 | -1.00 |  |  | -0.75 | -0.88 |  |  | -4.74 | -1.51 |  |  | -2.93 | -0.15 |  |  |  | |  | |  |
| GTDB 220.0 | 456.40 | 457.20 | 456.60 |  | 430.40^c^ | 436.00^b^ | 446.60^a^ |  | 338.00^c^ | 358.20^b^ | 375.20^a^ |  | 216.60^c^ | 243.60^b^ | 274.00^a^ |  | 0.2076 | < 0.0001 | | < 0.0001 | | < 0.0001 |
|  |  | 0.18 | 0.04 |  |  | 1.30 | 3.76 |  |  | 5.98 | 11.01 |  |  | 12.47 | 26.50 |  |  |  | |  | |  |
| NCBI 2024.10 | 423.20^b^ | 425.20^b^ | 437.00^a^ |  | 276.40^c^ | 284.40^b^ | 308.80^a^ |  | 177.00^c^ | 206.00^b^ | 225.60^a^ |  | 142.20^c^ | 158.00^b^ | 182.60^a^ |  | < 0.0001 | < 0.0001 | | < 0.0001 | | < 0.0001 |
|  |  | 0.47 | 3.26 |  |  | 2.89 | 11.72 |  |  | 16.38 | 27.46 |  |  | 11.11 | 28.41 |  |  |  | |  | |  |
| RDP  2.14 | 437.00^b^ | 448.00^a^ | 442.20^b^ |  | 304.60^b^ | 336.80^a^ | 344.60^a^ |  | 182.60^c^ | 227.40^b^ | 241.80^a^ |  | 134.20^c^ | 159.20^b^ | 183.40^a^ |  | 0.0062 | < 0.0001 | | < 0.0001 | | < 0.0001 |
|  |  | 2.52 | 1.19 |  |  | 10.57 | 13.13 |  |  | 24.53 | 32.42 |  |  | 18.63 | 36.66 |  |  |  | |  | |  |
| SILVA 138.1 | 458.20 | 458.20 | 458.20 |  | 453.80^b^ | 454.40^ab^ | 454.80^a^ |  | 442.80^a^ | 431.80^c^ | 437.80^b^ |  | 306.80^c^ | 321.20^b^ | 344.00^a^ |  | - | 0.0399 | | < 0.0001 | | < 0.0001 |
|  |  | 0.00 | 0.00 |  |  | 0.13 | 0.22 |  |  | -2.48 | -1.13 |  |  | 4.69 | 12.13 |  |  |  | |  | |  |
| SILVA 138.2 | 458.20 | 458.20 | 458.20 |  | 454.00^a^ | 450.80^b^ | 454.20^a^ |  | 438.80^a^ | 428.80^b^ | 437.60^a^ |  | 305.20^c^ | 323.00^b^ | 332.40^a^ |  | - | < 0.0001 | | < 0.0001 | | < 0.0001 |
|  |  | 0.00 | 0.00 |  |  | -0.70 | 0.04 |  |  | -2.28 | -0.27 |  |  | 5.83 | 8.91 |  |  |  | |  | |  |
| (2) Based on the V3-V4 amplicon sequences | | | | | | | | | | | | | | | | | | | | | | |
|  | Measurements (%) | | | | | | | | | | | | | | |  |  | |  | |  |  |
| Taxonomy | Phylum | | |  | Family | | |  | Genus | | |  | Species | | |  | *P*-value | | | | | |
| Classifier | UWTC | AWTC | MWTC |  | UWTC | AWTC | MWTC |  | UWTC | AWTC | MWTC |  | UWTC | AWTC | MWTC |  | P | | F | | G | S |
| GG2 2022.10 | 1464.00^a^ | 1430.50^b^ | 1332.75^c^ |  | 1366.00^a^ | 1329.50^b^ | 1287.25^c^ |  | 1087.75^a^ | 1023.75^b^ | 1070.25^a^ |  | 654.75^a^ | 611.75^b^ | 664.00^a^ |  | < 0.0001 | | < 0.0001 | | 0.0003 | 0.0006 |
|  |  | -2.29 | -8.97 |  |  | -2.67 | -5.77 |  |  | -5.88 | -1.61 |  |  | -6.57 | 1.41 |  |  | |  | |  |  |
| GG2 2024.09 | 1446.00^a^ | 1377.25^b^ | 1320.00^c^ |  | 1348.00^a^ | 1294.75^b^ | 1272.00^c^ |  | 1077.50^a^ | 999.75^b^ | 1060.75^a^ |  | 640.75^b^ | 566.00^c^ | 663.50^a^ |  | < 0.0001 | | < 0.0001 | | < 0.0001 | < 0.0001 |
|  |  | -4.75 | -8.71 |  |  | -3.95 | -5.64 |  |  | -7.22 | -1.55 |  |  | -11.67 | 3.55 |  |  | |  | |  |  |
| GTDB 220.0 | 1457.00^b^ | 1491.00^a^ | 1491.50^a^ |  | 1332.25^c^ | 1349.00^b^ | 1399.75^a^ |  | 993.00^b^ | 983.75^b^ | 1106.25^a^ |  | 520.25^c^ | 571.75^b^ | 697.25^a^ |  | < 0.0001 | | < 0.0001 | | < 0.0001 | < 0.0001 |
|  |  | 2.33 | 2.37 |  |  | 1.26 | 5.07 |  |  | -0.93 | 11.40 |  |  | 9.90 | 34.02 |  |  | |  | |  |  |
| NCBI 2024.10 | 1366.00^b^ | 1432.50^a^ | 1422.25^a^ |  | 821.75^c^ | 891.25^b^ | 952.75^a^ |  | 416.00^c^ | 583.00^b^ | 617.75^a^ |  | 266.25^c^ | 420.75^b^ | 485.25^a^ |  | < 0.0001 | | < 0.0001 | | < 0.0001 | < 0.0001 |
|  |  | 4.87 | 4.12 |  |  | 8.46 | 15.94 |  |  | 40.14 | 48.50 |  |  | 58.03 | 82.25 |  |  | |  | |  |  |
| RDP  2.14 | 1386.50^c^ | 1433.25^b^ | 1470.50^a^ |  | 888.00^c^ | 970.75^b^ | 1060.00^a^ |  | 390.75^c^ | 545.75^b^ | 624.00^a^ |  | 255.75^c^ | 365.25^b^ | 453.25^a^ |  | < 0.0001 | | < 0.0001 | | < 0.0001 | < 0.0001 |
|  |  | 3.37 | 6.06 |  |  | 9.32 | 19.37 |  |  | 39.67 | 59.69 |  |  | 42.82 | 77.22 |  |  | |  | |  |  |
| SILVA 138.1 | 1531.50^a^ | 1525.50^a^ | 1505.25^b^ |  | 1449.25^a^ | 1420.25^b^ | 1439.75^a^ |  | 1275.75^b^ | 1202.75^c^ | 1328.75^a^ |  | 603.50^c^ | 915.75^a^ | 860.25^b^ |  | 0.0036 | | 0.0081 | | 0.0002 | < 0.0001 |
|  |  | -0.39 | -1.71 |  |  | -2.00 | -0.66 |  |  | -5.72 | 4.15 |  |  | 51.74 | 42.54 |  |  | |  | |  |  |
| SILVA 138.2 | 1530.00^a^ | 1530.25^a^ | 1505.00^b^ |  | 1447.25^a^ | 1422.75^b^ | 1441.50^a^ |  | 1277.50^b^ | 1134.00^c^ | 1325.00^a^ |  | 617.00^c^ | 862.75^a^ | 808.00^b^ |  | 0.0021 | | 0.0122 | | < 0.0001 | < 0.0001 |
|  |  | 0.02 | -1.63 |  |  | -1.69 | -0.40 |  |  | -11.23 | 3.72 |  |  | 39.83 | 30.96 |  |  | |  | |  |  |
| ^a-c^ Different superscripts in a row indicate statistically significant differences (*p* < 0.05).  ^1)^ UWTC, unweighted taxonomy classifier; AWTC, average weighted taxonomy classifier; MWTC, manually weighted taxonomy classifier  ^2)^ P, phylum; F, family; G, genus; S, species | | | | | | | | | | | | | | | | | | | | | | |

| **Table S10.** Average values and statistical analysis results of fully-classified ratios for the *in vitro* dataset. The percentage difference was calculated to indicate how much AWTC and MWTC differed from UWTC. | | | | | | | | | | |
| --- | --- | --- | --- | --- | --- | --- | --- | --- | --- | --- |
| (1) Based on the full-length amplicon sequences | | | | | | | | | | |
|  | Measurements | | | | | | |  |  |  |
| Taxonomy level | Genus | | |  | Species | | |  | *P*-value | |
| Classifier ^1)^ | UWTC | AWTC | MWTC |  | UWTC | AWTC | MWTC |  | Genus | Species |
| GG2 2022.10 | 89.48^a^ | 86.11^c^ | 88.96^b^ |  | 69.35^a^ | 64.41^b^ | 69.46^a^ |  | < 0.0001 | < 0.0001 |
|  |  | -3.77 | -0.59 |  |  | -7.13 | 0.16 |  |  |  |
| GG2 2024.09 | 87.91^a^ | 84.97^c^ | 87.29^b^ |  | 68.24^a^ | 64.43^b^ | 68.62^a^ |  | < 0.0001 | < 0.0001 |
|  |  | -3.35 | -0.72 |  |  | -5.58 | 0.56 |  |  |  |
| GTDB 220.0 | 75.24^c^ | 84.37^b^ | 88.01^a^ |  | 48.44^c^ | 53.86^b^ | 61.52^a^ |  | < 0.0001 | < 0.0001 |
|  |  | 12.13 | 16.97 |  |  | 11.18 | 27.01 |  |  |  |
| NCBI 2024.10 | 42.32^c^ | 48.02^b^ | 50.45^a^ |  | 35.18^c^ | 37.44^b^ | 43.91^a^ |  | < 0.0001 | < 0.0001 |
|  |  | 13.46 | 19.21 |  |  | 6.42 | 24.80 |  |  |  |
| RDP 2.14 | 44.29^c^ | 51.01^b^ | 54.40^a^ |  | 34.51^c^ | 39.40^b^ | 44.32^a^ |  | < 0.0001 | < 0.0001 |
|  |  | 15.18 | 22.84 |  |  | 14.17 | 28.43 |  |  |  |
| SILVA 138.1 | 97.68^a^ | 96.70^c^ | 97.28^b^ |  | 74.78^b^ | 74.16^c^ | 80.83^a^ |  | < 0.0001 | < 0.0001 |
|  |  | -1.01 | -0.41 |  |  | -0.83 | 8.09 |  |  |  |
| SILVA 138.2 | 97.15^b^ | 96.27^c^ | 97.28^a^ |  | 75.60^b^ | 74.11^c^ | 79.44^a^ |  | < 0.0001 | < 0.0001 |
|  |  | -0.91 | 0.14 |  |  | -1.97 | 5.08 |  |  |  |
| (2) Based on the V3-V4 amplicon sequences | | | | | | | | | | |
|  | Measurements (%) | | | | | | |  |  |  |
| Taxonomy level | Genus | | |  | Species | | |  | *P*-value | |
| Classifier ^1)^ | UWTC | AWTC | MWTC |  | UWTC | AWTC | MWTC |  | Genus | Species |
| GG2 2022.10 | 81.27^a^ | 76.77^b^ | 80.78^a^ |  | 56.54^a^ | 51.26^c^ | 54.73^b^ |  | < 0.0001 | < 0.0001 |
|  |  | -5.54 | -0.60 |  |  | -9.35 | -3.21 |  |  |  |
| GG2 2024.09 | 81.39^a^ | 77.20^b^ | 80.79^a^ |  | 55.63^a^ | 50.32^b^ | 55.59^a^ |  | 0.0005 | < 0.0001 |
|  |  | -5.15 | -0.74 |  |  | -9.54 | -0.08 |  |  |  |
| GTDB 220.0 | 74.85^b^ | 73.64^b^ | 78.87^a^ |  | 38.03^c^ | 43.10^b^ | 50.13^a^ |  | 0.0005 | < 0.0001 |
|  |  | -1.62 | 5.37 |  |  | 13.32 | 31.81 |  |  |  |
| NCBI 2024.10 | 36.44^c^ | 46.99^b^ | 52.80^a^ |  | 25.77^c^ | 35.63^b^ | 42.42^a^ |  | < 0.0001 | < 0.0001 |
|  |  | 28.96 | 44.91 |  |  | 38.26 | 64.61 |  |  |  |
| RDP 2.14 | 34.87^c^ | 45.61^b^ | 53.42^a^ |  | 27.78^c^ | 34.29^b^ | 43.96^a^ |  | < 0.0001 | < 0.0001 |
|  |  | 30.80 | 53.21 |  |  | 23.44 | 58.25 |  |  |  |
| SILVA 138.1 | 88.83^b^ | 86.72^c^ | 91.01^a^ |  | 46.12^c^ | 65.87^a^ | 62.53^b^ |  | < 0.0001 | < 0.0001 |
|  |  | -2.37 | 2.45 |  |  | 42.82 | 35.57 |  |  |  |
| SILVA 138.2 | 89.87^b^ | 81.02^c^ | 91.49^a^ |  | 47.05^b^ | 60.10^a^ | 60.14^a^ |  | < 0.0001 | < 0.0001 |
|  |  | -9.84 | 1.81 |  |  | 27.72 | 27.81 |  |  |  |
| ^a-c^ Different superscripts in a row indicate statistically significant differences (*p* < 0.05).  ^1)^ UWTC, unweighted taxonomy classifier; AWTC, average weighted taxonomy classifier; MWTC, manually weighted taxonomy classifier | | | | | | | | | | |

| **Table S11.** Average values and statistical analysis results of error rates for both *in vivo* and *in vitro* datasets. The percentage difference was calculated to indicate how much AWTC and MWTC differed from UWTC. | | | | | | | | | | |
| --- | --- | --- | --- | --- | --- | --- | --- | --- | --- | --- |
| (1) Based on the *in vivo* dataset | | | | | | | | | | |
|  | Measurements | | | | | | |  |  |  |
| Sequences | Full-length amplicon | | |  | V3-V4 amplicon | | |  | *P*-value | |
| Classifier ^1)^ | UWTC | AWTC | MWTC |  | UWTC | AWTC | MWTC |  | Full length | V3-V4 |
| Phylum | 1.48^ab^ | 1.51^a^ | 1.34^b^ |  | 6.75^b^ | 6.31^c^ | 7.18^a^ |  | 0.0113 | < 0.0001 |
|  |  | 1.99 | -9.24 |  |  | -6.60 | 6.37 |  |  |  |
| Class | 5.15^a^ | 4.16^b^ | 4.15^b^ |  | 17.03^a^ | 13.70^c^ | 14.74^b^ |  | < 0.0001 | < 0.0001 |
|  |  | -19.15 | -19.48 |  |  | -19.57 | -13.43 |  |  |  |
| Order | 11.80^a^ | 9.93^b^ | 10.18^b^ |  | 26.92^a^ | 24.80^b^ | 24.07^c^ |  | < 0.0001 | < 0.0001 |
|  |  | -15.85 | -13.77 |  |  | -7.85 | -10.56 |  |  |  |
| Family | 37.06^a^ | 36.23^a^ | 35.46^b^ |  | 39.23^a^ | 39.30^a^ | 36.68^b^ |  | < 0.0001 | < 0.0001 |
|  |  | -2.23 | -4.31 |  |  | 0.17 | -6.51 |  |  |  |
| Genus | 55.00^b^ | 58.60^a^ | 50.29^c^ |  | 70.92^b^ | 71.80^a^ | 67.39^c^ |  | < 0.0001 | < 0.0001 |
|  |  | 6.55 | -8.57 |  |  | 1.24 | -4.97 |  |  |  |
| Species | 82.96^a^ | 79.73^c^ | 79.83^b^ |  | 90.50^a^ | 90.79^a^ | 87.08^b^ |  | < 0.0001 | < 0.0001 |
|  |  | -3.89 | -3.78 |  |  | 0.32 | -3.79 |  |  |  |
| (2) Based on the *in vitro* dataset | | | | | | | | | | |
|  | Measurements | | | | | | |  |  |  |
| Sequences | Full-length amplicon | | |  | V3-V4 amplicon | | |  | *P*-value | |
| Classifier ^1)^ | UWTC | AWTC | MWTC |  | UWTC | AWTC | MWTC |  | Full length | V3-V4 |
| Phylum | 4.98 | 4.76 | 4.45 |  | 12.91 | 12.20 | 13.00 |  | 0.1839 | 0.3247 |
|  |  | -4.32 | -10.56 |  |  | -5.46 | 0.74 |  |  |  |
| Class | 8.76^a^ | 8.06^ab^ | 7.86^b^ |  | 24.95^a^ | 21.32^b^ | 22.09^b^ |  | 0.0149 | 0.0017 |
|  |  | -8.01 | -10.35 |  |  | -14.54 | -11.46 |  |  |  |
| Order | 24.17^a^ | 22.62^b^ | 21.68^b^ |  | 36.64^a^ | 34.07^b^ | 33.35^b^ |  | 0.0002 | 0.0011 |
|  |  | -6.42 | -10.33 |  |  | -7.03 | -8.97 |  |  |  |
| Family | 46.12^a^ | 45.47^a^ | 42.77^b^ |  | 52.29^a^ | 52.01^a^ | 49.67^b^ |  | < 0.0001 | 0.0001 |
|  |  | -1.42 | -7.27 |  |  | -0.55 | -5.02 |  |  |  |
| Genus | 68.95^a^ | 69.29^a^ | 64.75^b^ |  | 79.22^a^ | 79.62^a^ | 75.68^b^ |  | < 0.0001 | < 0.0001 |
|  |  | 0.50 | -6.10 |  |  | 0.50 | -4.48 |  |  |  |
| Species | 80.46^a^ | 80.73^a^ | 76.07^b^ |  | 91.29^a^ | 91.47^a^ | 87.94^b^ |  | < 0.0001 | < 0.0001 |
|  |  | 0.33 | -5.46 |  |  | 0.20 | -3.67 |  |  |  |
| ^a-c^ Different superscripts in a row indicate statistically significant differences (*p* < 0.05).  ^1)^ UWTC, unweighted taxonomy classifier; AWTC, average weighted taxonomy classifier; MWTC, manually weighted taxonomy classifier. | | | | | | | | | | |
