## Supplementary Figures S1 to S6 for "Manually weighted taxonomy classifiers improve species-specific rumen microbiome analysis compared to unweighted or average weighted taxonomy classifiers"

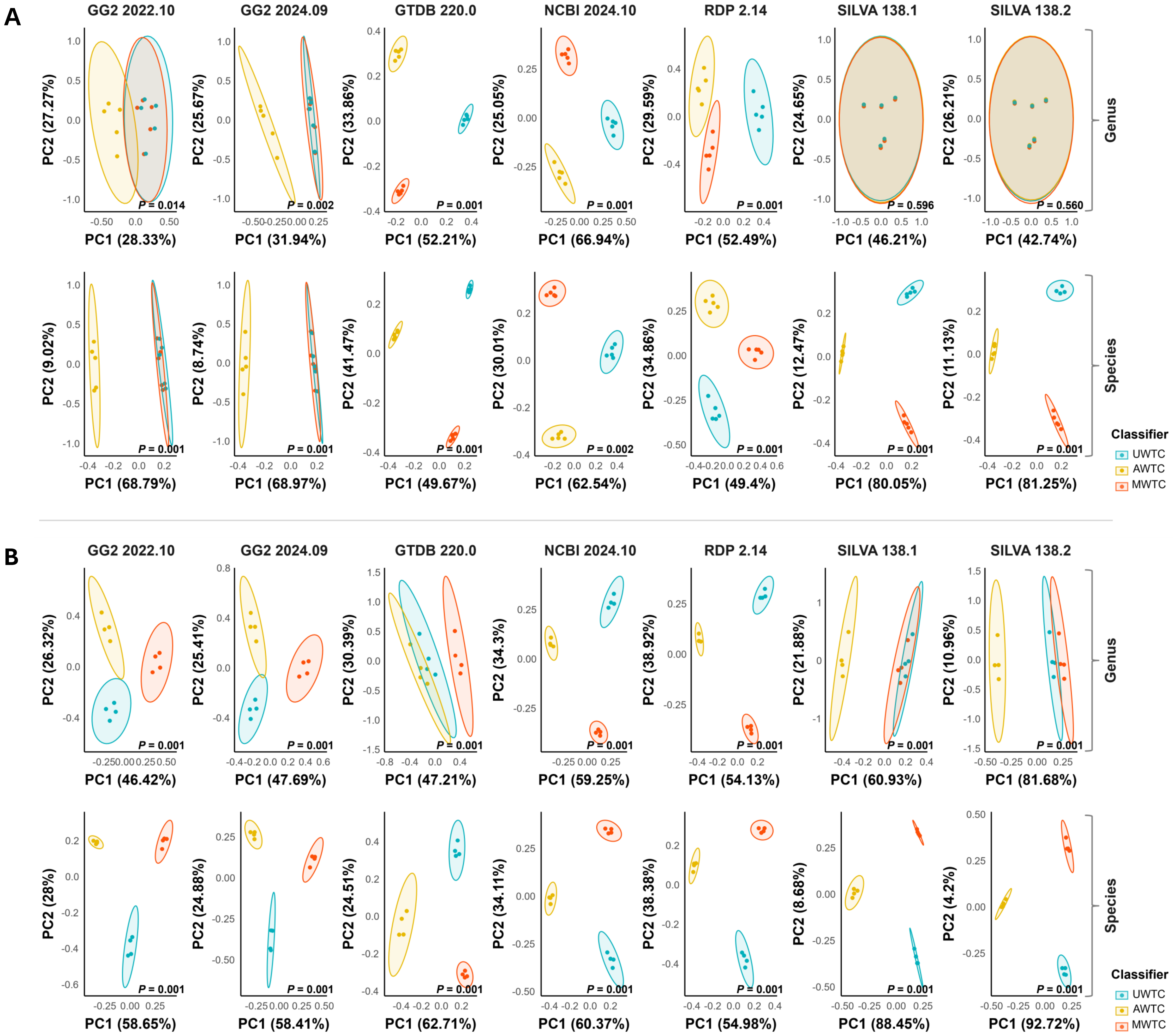


**Fig. S1.** PCA plots showing differences in rumen microbiome between classifier types within each database at the genus and the species levels for the full-length amplicon sequences (A), and at the genus and the species levels for the V3-V4 amplicon sequences (B) in an *in vitro* study. Significant differences were observed across all the classifiers except at the genus level with the SILVA database.

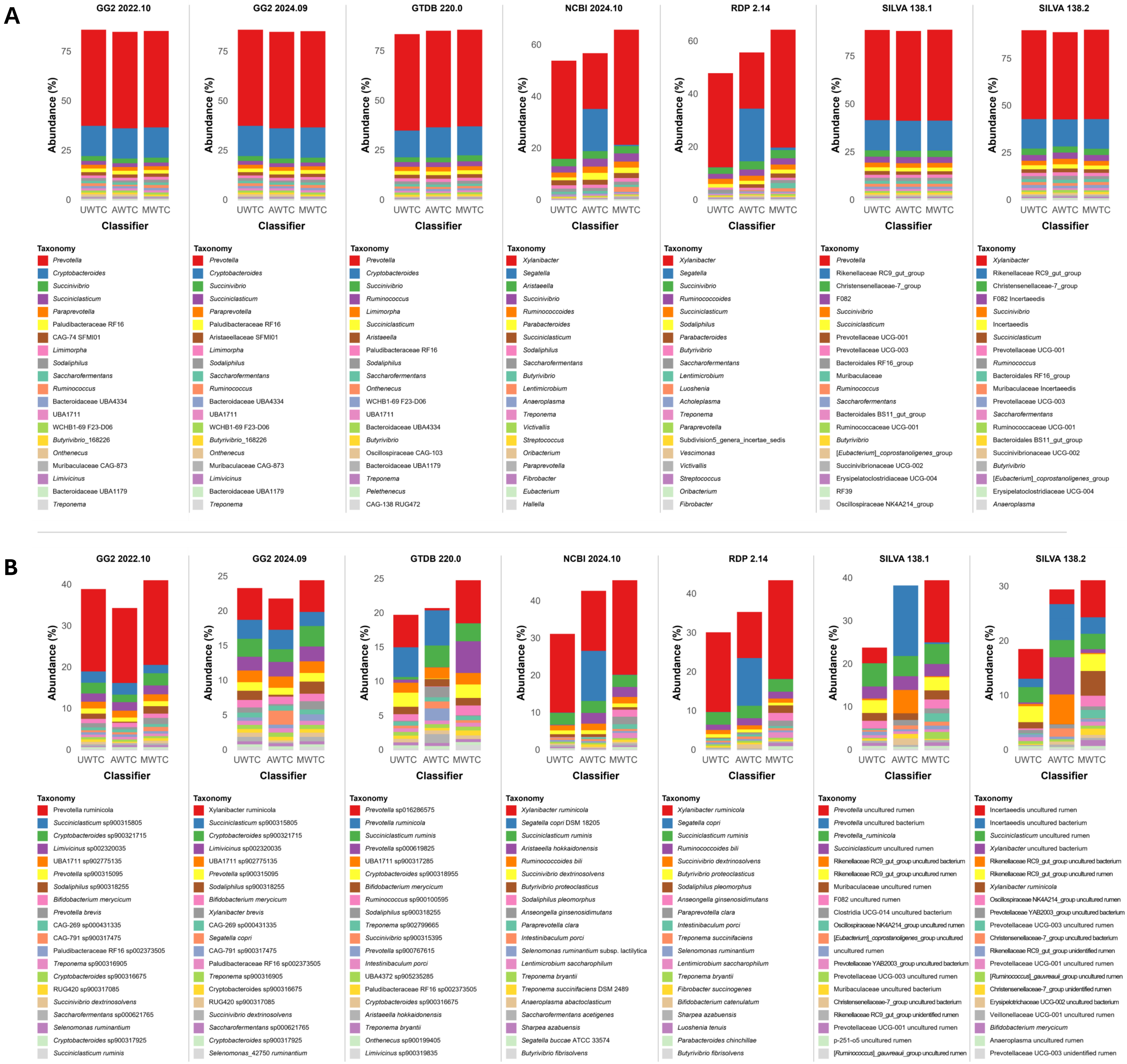


**Fig. S2.** Taxonomy barplot showing the top 20 taxa for each taxonomy classifier in an *in vivo* study. Taxonomy classification was performed at the genus level using full-length amplicon sequences (A) and at the species level using V3-V4 amplicon sequences (B).

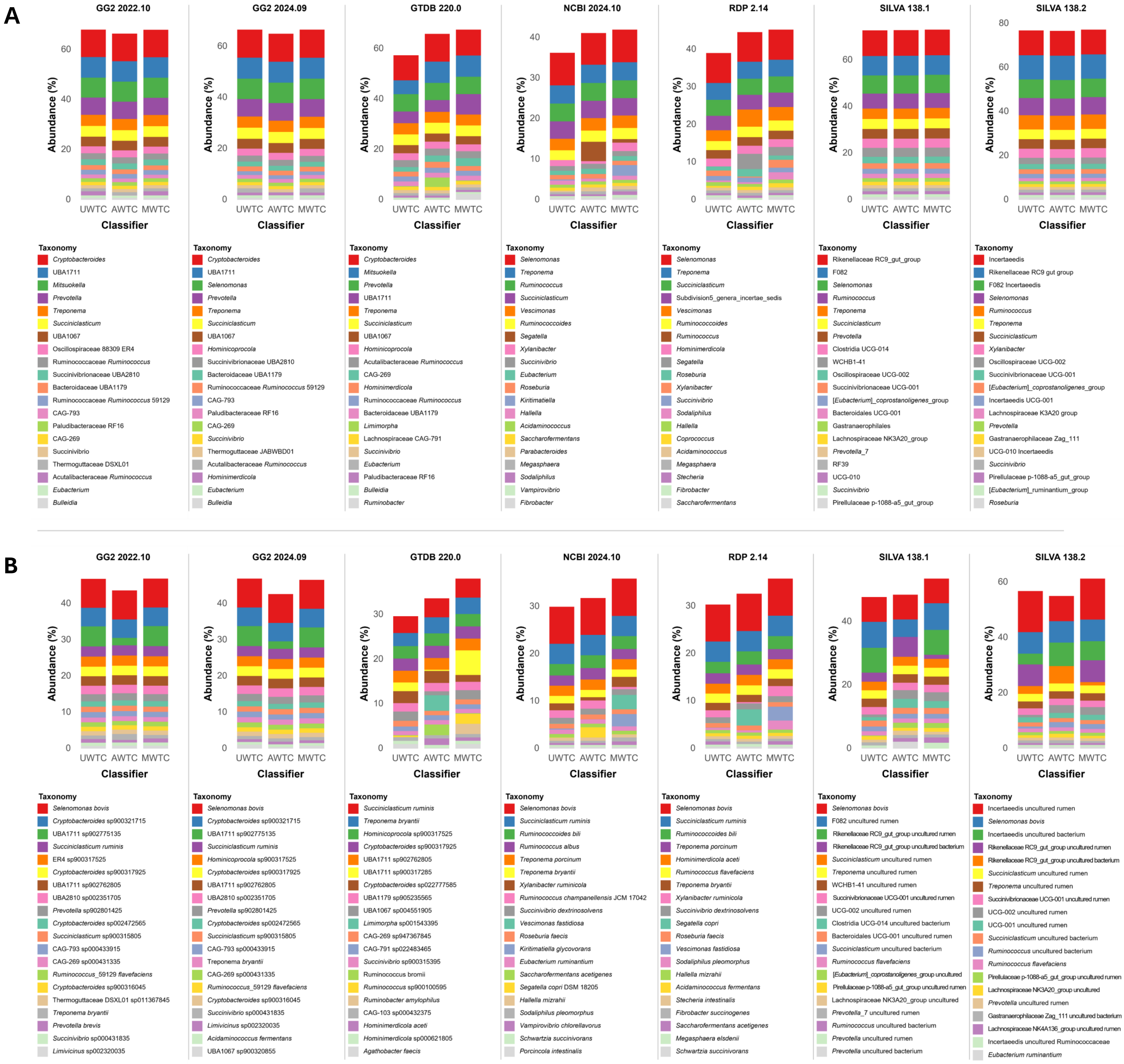


**Fig. S3.** Taxonomic barplot showing the top 20 taxa for each taxonomy classifier in an *in vitro* study. Taxonomy classification was performed using full-length amplicon sequences at the genus (A) and species level (B).

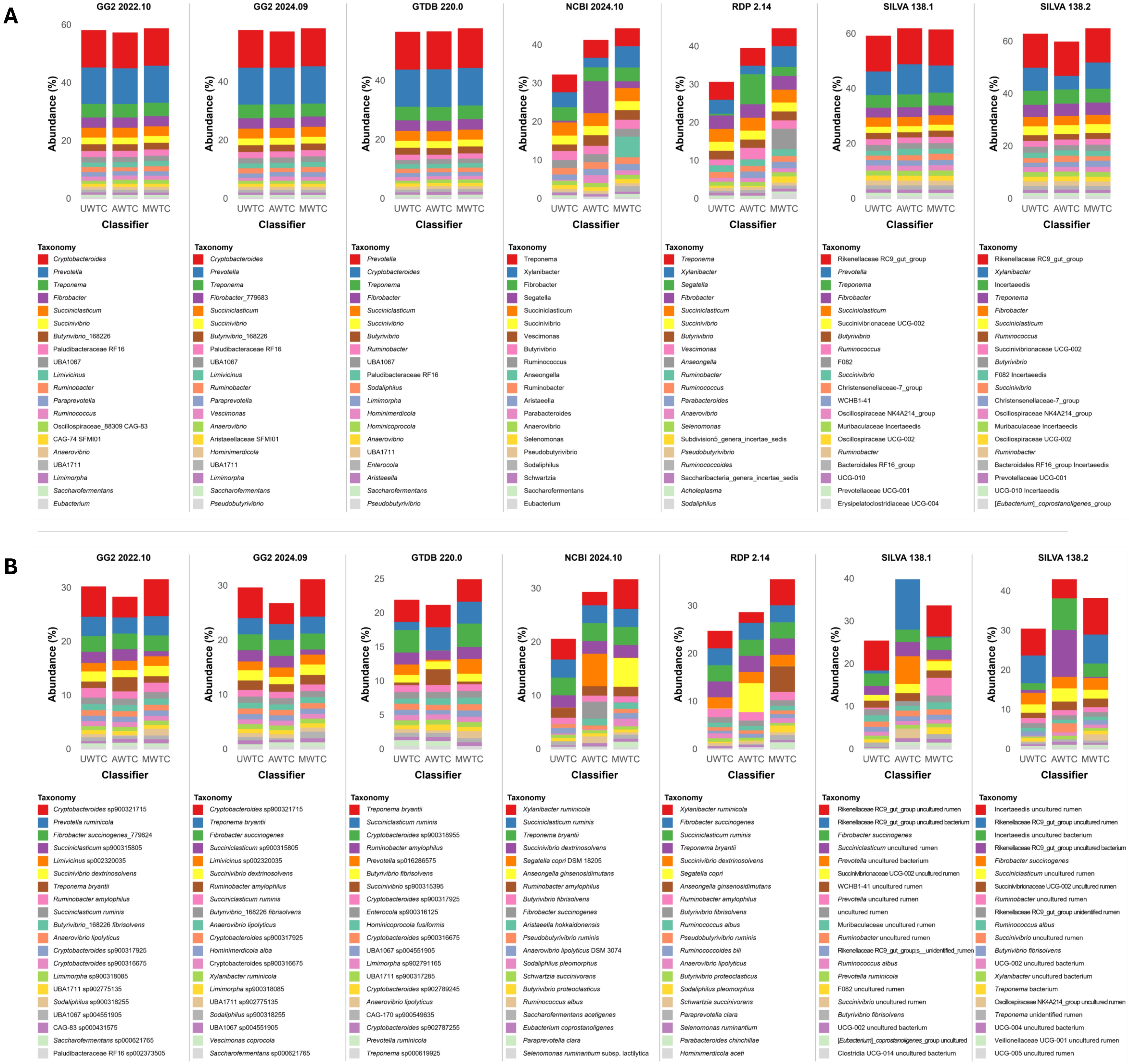


**Fig. S4.** Taxonomic barplot showing the top 20 taxa for each taxonomy classifier in an *in vitro* study. Taxonomy classification was performed using V3-V4 amplicon sequences at the genus (A) and species level (B).

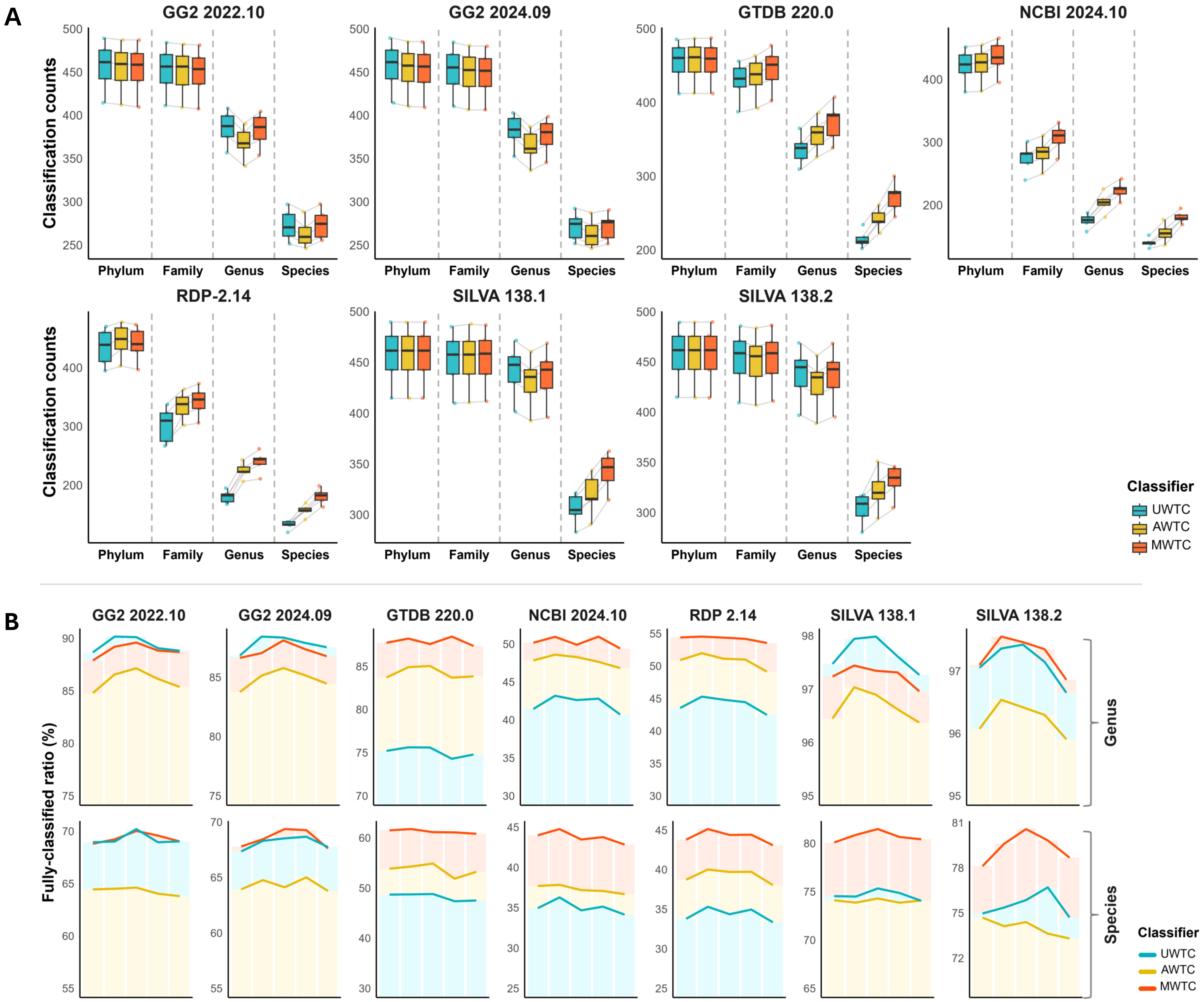


**Fig. S5.** Counts of classified ASVs at the phylum, family, genus, and species levels for each database (A) and the fully classified ratios (the proportion of completely classified features relative to the total ASVs) (B) for each database identified using the full-length amplicon sequences in an in vitro study. Classifiers were categorized as follows: UWTC, a classifier without any adjustments; AWTC, a classifier based on average data from the EMPO3 dataset; and MWTC, a classifier manually curated using metagenomic and amplicon sequencing data.

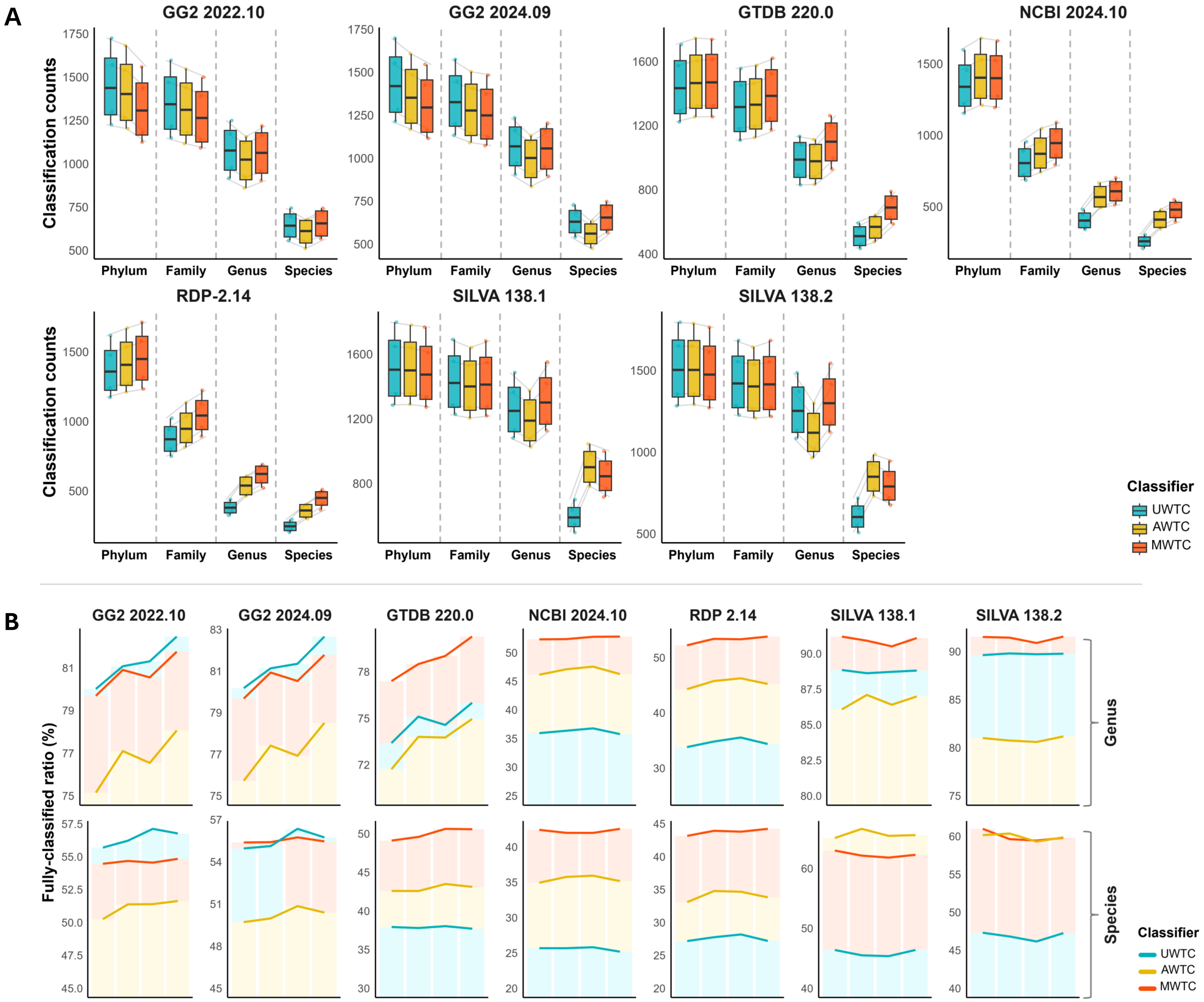


**Fig. S6.** Counts of classified ASVs at the phylum, family, genus, and species levels for each database (A) and the fully classified ratios (the proportion of completely classified features relative to the total ASVs) (B) for each database identified using the V3-V4 amplicon sequences in an in vitro study. Classifiers were categorized as follows: UWTC, a classifier without any adjustments; AWTC, a classifier based on average data from the EMPO3 dataset; and MWTC, a classifier manually curated using metagenomic and amplicon sequencing data.
